# Exercise-responsive extracellular vesicles-associated miR-196a-5p and miR-342-3p as candidate mediators of skeletal muscle-adipose tissue crosstalk

**DOI:** 10.1101/2025.05.28.656129

**Authors:** Dominik Tischer, Daniela Schranner, Sebastian Kallabis, Alexander Braunsperger, Martin Schönfelder, Thorsten Gnad, Paul Jonas Jost, Svetozar Nesic, Florian Renziehausen, Lars Fester, Andreas Buness, Jan Hasenauer, Felix Meissner, Alexander Pfeifer, Henning Wackerhage, Ana Soriano-Arroquia

## Abstract

Despite evidence showing that exercise triggers the expression of thermogenic markers in murine models, it is still an ongoing debate whether similar mechanisms occur in humans. The aim of this study was to investigate the molecular mechanisms that might mediate the crosstalk between thermogenic adipose tissue and skeletal muscle during exercise in humans. To do so, we isolated the serum from sedentary individuals, bodybuilders, sprinters and endurance athletes before and after a maximal graded exercise test. Our results show that post-exercise serum from endurance athletes increases *UCP1* transcription in human white adipocytes. Moreover, treatment of human white adipocytes with muscle-derived extracellular vesicles (EVs) increased the expression of *UCP1*. The levels of *UCP1* were boosted when white adipocytes were treated with EVs from muscle cells after electrical pulse stimulation (EPS) compared to non-stimulated muscle EVs. Small-RNA-sequencing of serum-derived EVs from endurance athletes before and after exercise and systematic database collection based on PRISMA guidelines revealed miR-196a/b-5p and miR-342-3p as promising exercise-induced candidate EV-associated microRNAs to investigate. Particularly, we found that muscle cells release EVs enriched in miR-196a-5p after EPS and cAMP treatment, a microRNA that increases *UCP1* expression in human white adipocytes. Likewise, miR-342-3p is released from human white and brown adipocytes EVs upon cAMP stimulation. Overexpression of miR-342-3p increased *MYOG* and *PGC1α* transcriptional levels during myogenic differentiation. Our results suggest that a bout of exercise to exhaustion in endurance athletes releases circulatory EVs enriched in microRNAs that regulate the expression of genes involved in browning and muscle differentiation.

**Graphical abstract:** 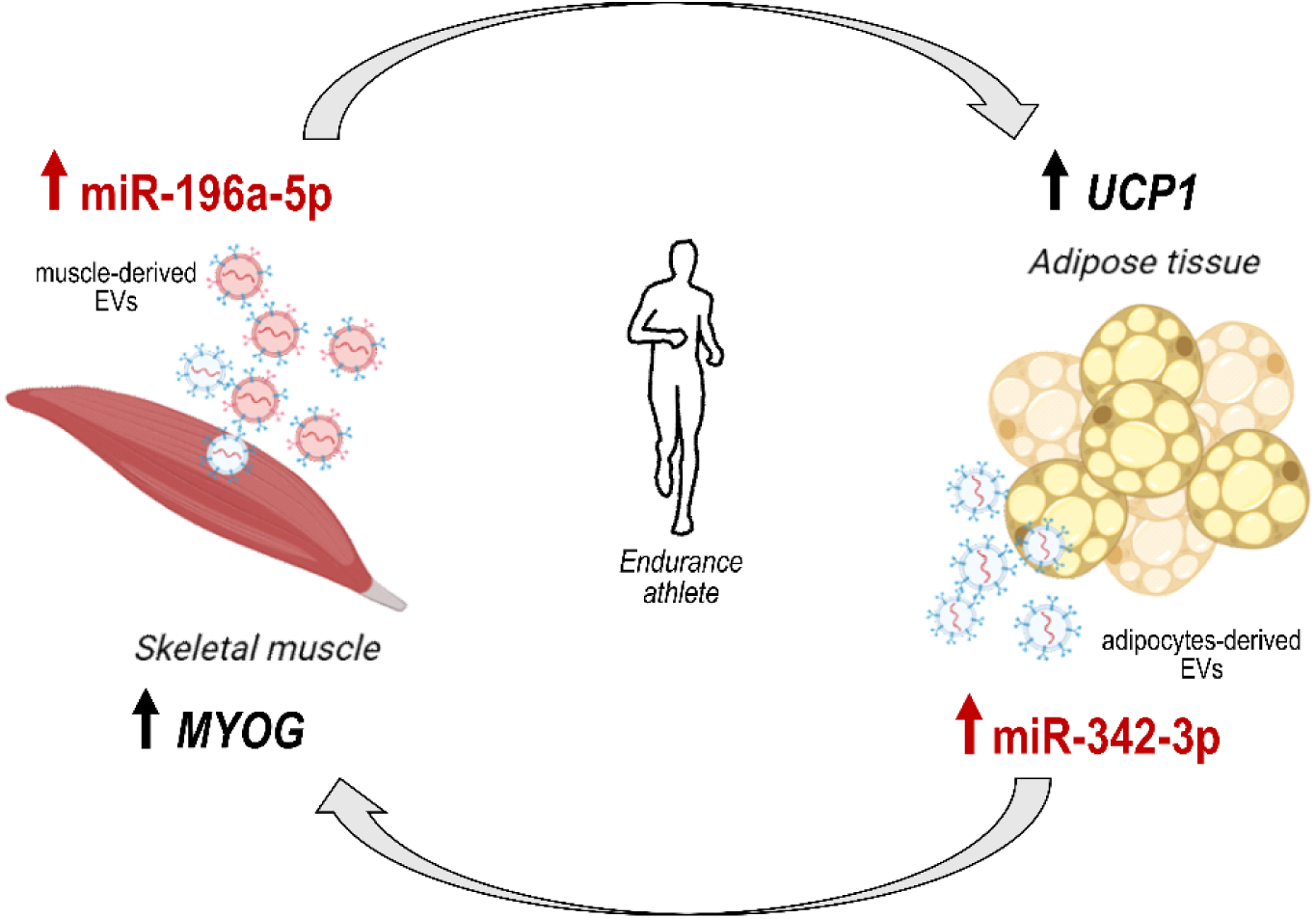

**Highlights:**

- Post-exercise serum from endurance athletes increases *UCP1* transcriptional levels in human white adipocytes.
- Muscle-derived EVs induce *UCP1* expression in human white adipocytes.
- miR-196a and miR-342-3p are increased in serum-EVs from endurance athletes after a maximum graded bicycle test to exhaustion.
- Muscle cells release EVs enriched in miR-196a after electrical pulse stimulation.
- Brown and white adipocytes release EVs enriched in miR-342-3p after cAMP stimulation.
- miR-196a and miR-342-3p have browning and myogenic potential, respectively.

## Introduction

Extracellular vesicles (EVs) are lipid bilayer particles without a nucleus that lack the capacity to replicate on their own (Welsh *et al*., 2024). EVs can be classified into different categories depending on their biogenesis and size. Small EVs usually refer to those EVs with a size of <200 nm in diameter, sub-classified into exosomes, which are membrane-enclosed vesicles of endocytic origin released by the multivesicular body (MVB); and ectosomes (also known as microvesicles), which refer to plasma-derived vesicles (Welsh *et al*., 2024). Originally thought of as cellular garbage, EVs are nowadays known for their biological function and role in cell-to-cell communication, as they carry active molecules such as microRNAs that are taken up by neighbouring or long-distance cells (Valadi *et al*., 2007; Vechetti *et al*., 2021). Due to their lipid bilayer, EVs prevent RNA degradation through ribonucleases present in the blood, resulting in an efficient way of transporting microRNAs between cells and tissue. Therefore, microRNAs in EVs are a better source for transcriptomic analyses compared to free circulating microRNAs (Cheng *et al*., 2014). microRNAs are small non-coding RNAs of approximately 22 nucleotides that regulate gene expression post-transcriptionally (Lee *et al*., 1993; Wightman *et al*., 1993). They inhibit gene expression, especially through mRNA decay or directly repressing protein translation by binding to partially complementary sites of their target transcripts at the 3’ untranslated region (3’UTR), although interactions at the 5’UTR or coding regions have also been described (Iwakawa & Tomari, 2015; O’Brien *et al*., 2018).

MicroRNAs contained in small EVs, here referred to as EV-associated microRNAs, are released into the extracellular space and circulation by cells under certain stimuli such as exercise (Valadi *et al*., 2007). Advances in high-throughput omics and EV characterization made it possible to explore new ways of communication between distant tissues through EVs and microRNAs. For example, muscle-to-adipose tissue interactions involve the release of myokines such as IL-6 and irisin, some of which have been detected in circulating EVs after exercise, suggested to induce browning or beiging of white adipose tissue (Severinsen & Pedersen, 2020; Shi *et al*., 2024). Unlike white adipose tissue, brown and beige adipose tissue contribute to non-shivering thermogenesis by releasing energy in the form of heat during oxidative phosphorylation at the expense of ATP production, resulting in increased energy expenditure. This process is mediated by the mitochondrial protein uncoupling protein 1 (UCP1), which is triggered especially via adrenergic signalling during cold exposure and diet-induced thermogenesis (Cypess *et al*., 2015; U Din *et al*., 2018). Induced thermogenesis through white-to-beige conversion of white adipose tissue, a process known as ‘browning’ or ‘beiging’, has been proposed as a potent therapeutic way for combating metabolic diseases and obesity (Wu *et al*., 2014).

Despite previous data showing that exercise increases UCP1 levels in subcutaneous white adipose tissue and increases whole-body energy expenditure in rats (Wu *et al*., 2014), discrepancies between rodent and human research and conflicting results still question whether exercise contributes to white adipose tissue browning in humans (Otero-Díaz *et al*., 2018; Tsiloulis *et al*., 2018; Severinsen & Pedersen, 2020; Braunsperger *et al*., 2026). Additionally, little is known about possible exercise-induced adipose-to-muscle interactions. Interestingly, it has been previously shown that a bout of incremental exercise at the anaerobic threshold induces a greater response of circulating myokines and adipokines such as FGF-21, follistatin and omentin compared to aerobic exercise performed at maximum fat oxidation rate (He *et al*., 2019). Notably, it has been also suggested that a bigger effect is needed to induce thermogenic markers in white adipose tissue after exercise in humans (Otero-Díaz *et al*., 2018). The aim of this study was to screen for exercise-regulated microRNAs mediating bi-directional crosstalk between skeletal muscle and thermogenic adipose tissue. Here, we identify exercise-regulated EV-associated microRNAs that may contribute to skeletal muscle-adipose tissue crosstalk. We show that a bout of maximal graded exercise test in endurance athletes promotes the release of circulating microRNAs such as miR-196a-5p and miR-342-3p that regulate the expression of genes involved in browning and muscle differentiation, respectively. Our findings reveal new insights about the intricate relationship between both tissues during exercise in humans.

## Methods

### Ethical approval and human samples collection

This study conforms to the Declaration of Helsinki for the use of human subjects and tissues, and it was approved by the medical ethics committee of the Technical University of Munich (356/17S). Exact details of the human exercise intervention and serum sample collection can be found in Schranner et al., Physiol Rep. (2021) (Schranner *et al*., 2021). Briefly, all participants performed the same exercise test, consisted on a ramp-test on a bicycle ergometer with power increasing linearly (30 W / minute) until voluntary exhaustion. A exhaustion criteria of either a respiratory exchange ratio (RER) >1.0 or a ventilatory equivalent of oxygen (VEeqO_2_) of >30.0 was used, ensuring that each participant achieved the maximal oxygen uptake (VO_2_max) to determine endurance performance. Blood samples were obtained in the fasted state from the antecubital vein before and 5 minutes after the graded exercise test to exhaustion in healthy male participants. Participants were either: sedentary controls, endurance athletes, natural bodybuilders or sprinters (see table below). For blood sampling, 9 ml serum S-Monovettes Z-Gel collecting tubes (Sarstedt AG und Co KG, Nuembrecht, Germany) was used. Blood samples were allowed to clot at room temperature for 30 minutes in an upright position. After centrifugation (10 min / 18°C, 2460 g), serum replicates were merged into one 15 ml Falcon tube (Greiner Bio-One GmbH, Kremsmuenster, Austria). Then, serum was aliquoted in cryotubes (Sarstedt AG und Co KG, Nümbrecht, Germany), frozen on dry ice for 30 min and stored at −80°C until analysis. Samples were not subjected to freeze-thaw cycles before EV isolation.

**Table 1.** Physical characteristics of sedentary controls, endurance athletes, natural bodybuilders and sprinters. Data is represented as mean ± SD. Abbreviations: BMI = body mass index; VT1 = Ventilatory threshold 1. Abs = absolute; Rel. = relative.

|  | **n** | **Age (years)** | **Weight (kg)** | **BMI (kg/m2)** | **Biceps circumference (cm)** | **Quadriceps circumference (cm)** | **VO<sub>2</sub>Max (ml/kg/min)** | **VT1 (ml/kg/min)** | **Abs. leg strength (Nm)** | **Rel.e leg strength (Nm/kg)** | **Hand grip strength (kg)** | **Reactive strength (RSI)** |
|---|---|---|---|---|---|---|---|---|---|---|---|---|
| Sedentary controls | 6 | 27 ± 3 | 81.5 ± 11.9 | 24.1 ± 2.6 | 29.8 ± 2.5 | 52.8 ± 5.4 | 43.9 ± 5.0 | 26.1 ± 1.9 | 321.2 ± 53.4 | 3.8 ± 0.4 | 58.7 ± 12.8 | 11.9 ± 2.9 |
| Endurance athletes (hWAs treatment) | 4 | 28 ± 2 | 72.5 ± 7.2 | 22.4 ± 0.8 | 28.3 ± 0.5 | 54.9 ± 2.4 | 64.8 ± 5.5 | 46.3 ± 5.5 | 449.3 ± 25.8 | 6.2 ± 0.3 | 52.8 ± 3.4 | 14.8 ± 5.7 |
| Endurance athletes (small RNA-seq) | 8 | 30 ± 5 | 74.7 ± 7.7 | 22.6 ± 2.1 | 29.3 ± 2.7 | 53.8 ± 5.9 | 63.5 ± 5.3 | 46.8 ± 6.4 | 431.6 ± 42.6 | 5.8 ± 0.3 | 55.8 ± 5.9 | 14.3 ± 2.5 |
| Natural bodybuilders | 6 | 30 ± 5 | 89.3 ± 7.1 | 28.0 ± 2 | 36.4 ± 2.8 | 61.5 ± 4.1 | 41.6 ± 3.5 | 25.2 ± 2.6 | 326.0 ± 32.6 | 3.7 ± 0.2 | 69.9 ± 5.3 | 13.0 ± 4.3 |
| Sprinters | 6 | 22 ± 3 | 76.1 ± 10.7 | 21.6 ± 1.4 | 28.5 ± 1.8 | 54.9 ± 4.1 | 52.3 ± 4.8 | 29.6 ± 3.3 | 359.7 ± 68.5 | 4.7 ± 0.3 | 58.8 ± 6.1 | 20.2 ± 3.3 |

### Cell culture and treatment of human white adipocytes with serum and small EVs

Cryopreserved subcutaneous human white adipocytes (hWAs) were purchased at passage 2 from Zenbio (SP-F-2; BMI 25.0-29.9). Cells were seeded, grown and expanded using Seeding Medium: DMEM/F-12 GlutaMAX (ThermoFisher, Cat. n. 31331-028); 15% Foetal Bovine Serum (FBS, Gibco, Cat. n. 10270-106); 7.5% horse serum (HS, Sigma-Aldrich, Cat. n. H1270); 0.7 mM L-glutamine (Sigma-Aldrich, Cat. n. G7513); 50,000 units of Penicillin and 50 mg of Streptomycin (P/S, Sigma-Aldrich, Cat. n. P0781); 1.25 µg of basic recombinant human Fibroblast Growth Factor (bFGF, PeproTech, Cat. n. 100-18B). For the experiments, a minimum of 25,000 cells/well in 24-well plates or 50,000 cells/well in 12-well plates were seeded at passage 4 and allowed to achieve over-confluency. When bright spots of pre-adipocytes clusters were clearly visible under the microscope, the medium was change to Zenbio pre-adipocytes Differentiation Medium (DM-2-500) and/or Maintenance Medium (AM-1) according to the manufacture’s guidelines.

For the treatment with small EVs, fully differentiated mature hWAswere washed twice with serum-free Treatment Medium (DMEM/F12 GlutaMax, ThermoFisher, Cat. n. 31331-028, without added components, without P/S). Fresh Treatment Medium was added with or without the treatment using a total volume of 300 µl/well or 600 µl/well for 24-well plates or 12-well plates, respectively. For the experiments, mature hWAs were treated with 10% v/v of serum or 7.8×10^9^ of concentrated serum-derived EVs per well for 16 hours in 24-well plates. Calculation for the treatment with serum-EVs was normalized based on the sample with the lowest EV amount according to the NTA measurement. Treatment of hWAs was performed simultaneously on the same day using the serum and serum-EVs from each individual to decrease technical variability. Paired-groups showing undetermined *UCP1* Ct values in either pre- or post-treated hWAs were excluded for the final analysis.

For the treatment of hWAs with non-stimulated and EPS-stimulated muscle-derived EVs, 1ug of purified small EVs (corresponding to 7.23×10^7^ of EVs) or an equal amount of PBS as non-EV treatment control were added per well in 12-well plates. This concentration was based on the highest respond to *UCP1* mRNA levels in hWAs after treatment titration optimization. EVs were dissolved in the same PBS buffer used as vehicle control (pH 7.4, #10010056, ThermoFisher). Cells were collected the next day 16 hours post-treatment for RNA isolation and Real-Time qPCR quantification.

### Cell isolation and differentiation of human brown adipocytes

Human surgeries for the isolation of brown adipose tissue were approved by the ethics committee of the Medical Faculty of the University of Bonn (Ethical registration numbers: No. 076/18) and North Rhine Medical Association (No. 2023214). All patients provided written informed consent before the surgery. Human BAT biopsies from the supraclavicular neck region were taken by a plastic surgeon. Biopsies were obtained during elective thyroid surgery at Johanniter Hospital Bonn. Cell isolation was performed according to previous published protocol (Gnad *et al*., 2020). Briefly, 500.000 cells from the Stromal Vascular Fraction (SVF) were seeded onto a 12-well plate in hBA growth media (GM): MEM/F-12, HEPES, no phenol red, (ThermoFisher, Cat. n. 11039-021) supplemented with 10% FBS (Sigma, Cat. n. F7524), 1% P/S (ThermoFisher, Cat. n. P06-07100) and 10µM FGF1 (Immuno Tools, Cat. n. 11343553). Growth media was replaced every second day. Once over-confluency was achieved, cells were induced for 3 days by replacing the media to hBA differentiation media (DM): MEM/F-12, HEPES, no phenol red, (ThermoFisher, Cat. n. 11039-021) supplemented with 1% P/S, 430nM insulin (Sigma, Cat. n. I9278), 1 µM dexamethason (Sigma, Cat. n. D4902), 1 nM triiodothyronine (Sigma, Cat. n. T6397), 10µg/ml transferrin (Lonza, Cat. n. CC-4205), 0.5 mM IBMX (Sigma, Cat. n. I5879) and 1 µM rosiglitazone (Sigma, Cat. n. R2408). After induction, media without IBMX but with 0.2 µM rosiglitazone was replaced for another 3 days. At day 6 post induction, differentiation media without IBMX or rosiglitazone was replaced. Media was changed every 3 days. Mature brown adipocytes were used for experiments at day 12 post-induction.

### EVs processing and visualization by Transmission Electron Microscopy (TEM)

Processing and embedding of EVs for TEM visualization was performed at room temperature on the same day after isolation and without freeze/thaw cycles. All solutions were filtered by 0.22 µm filters (Merck, Darmstadt, Germany). All TEM single slot grids were obtained from ScienceServices (Munich) coated by formvar membrane. Grids were prepared per protocol and air dried with sodium hydroxide pellets to prevent carbon dioxide exposure.

After isolation, EVs were fixed in 2 % glutaraldehyde, stored on ice and gently mixed in the reaction tube. For embedding, 2 % of Ultrapure L.M.P. agarose (Invitrogen) in aqua dest. were prepared in a microwave and stored for 15 minutes in an Eppendorf heater at 37°C. 50 µl L.M.P. Agarose were pipetted in a pre-warmed 2 ml reaction tube and incubated at 37°C. 5 µl of fixed EVs were added by mixing carefully in the viscous L.M.P. agarose and immediately stored on ice for solidification.

The solidified L.M.P. agarose was trimmed in pieces with 1 mm edge length and quickly stained with 2% O_2_O_4_ in for one hour at room temperature. Subsequently, samples were treated twice for 5 min. in ethanol with Phosphate Buffer in rising concentration (30%, 50%, 70%, 90%) and absolute ethanol three times for 5 minutes. The probes were carefully transferred in propylene oxide for 15 min twice at room temperature, followed by SPURR 1 hour two times changing in a vacuum chamber at 25°C and at final step at 70°C in a high vacuum for 24 hours. Probes were trimmed for ultrathin sections between 50 to 70 nm with an RMC PowerTome PC and dried in vacuum chamber for 4 hours. Contrasting with heavy metals was proceed using a Leica EM AC 20, with Uranylacetat in a concentration of 0.5 %, followed by 3 % lead citrate solution (Leica Ultrostain II). Subsequently, the probes on the Grids were carefully washed with carbon dioxide-free double distilled water and dried in a vacuum chamber surrounded with sodium hydroxide pellets. EVs were visualized using a TEM JEOL1400plus with tomography holder for high magnification, with 20.000x and 50.000x by TEM-Viewer, SightX-Viewer software (JEOL) and ImageJ Fiji adding marks (Schindelin *et al*., 2012).

### EV characterization via ZetaView Nanoparticle Tracking Analysis (NTA)

Extracellular vesicles concentration and size distribution were assessed using the ZetaView nanoparticle tracking analyzer (Particle Metrix, Germany). Prior to measurement, EV samples were diluted at a 1:500 ratio in filtered phosphate-buffered saline (PBS) to ensure optimal particle concentration within the instrument’s detection range. Measurements were performed in triplicate to ensure reproducibility. Sensitivity was adjusted to 80.0 to ensure optimal signal acquisition, and the shutter was set to 100 to regulate exposure effectively.

### PKH67 staining of EVs

Fluorescent staining of purified EVs was performed using PKH67 Green Fluorescent Mini Kit for General Cell Membrane Labeling (Sigma-Aldrich, #P7333) and according to the manufacture’s protocol. Briefly, 2ug of EVs were stained with 2X of Dye solution, immediately mixed by gentle pipetting for 30 seconds and incubated at room temperature for 10 minutes. Mock-stained control using 100% ethanol rather than the dye, with EVs, was used as the staining negative control. After incubation, 1 mL of EV-depleted horse serum (Sigma-Aldrich, Cat. n. H1270) by serial ultracentrifugation was added to each sample to stop the staining. Samples were centrifuged on 3-10 kDa cut-off weight filter columns (Amicon, #UFC200324) at 7500 xg for 15 minutes at 4°C. EVs were next concentrated in the concentrate collection tube by separating and inverting the filter device and spinning them down for 2 minutes at 1000 xg at 4°C. EVs were resuspended in PBS, placed on ice and immediately used for cell treatment. hWAs were incubated with the labelled EVs for 24 hours and fluorescent pictures were taken one day post-treatment.

### ExoView analysis of small EVs

Human serum-derived extracellular vesicles were diluted in manufacturer supplied incubation solution incubated overnight on ExoView Human Tetraspanin Plasma Chips (Unchained labs, US) featuring CD9 (clone HI9a), CD63 (clone H5C6), CD81(clone JS-81), CD41a (clone HIP8) and MIgG control (clone MOPC-21) capture antibodies following manufacturer’s instructions. Chips were then processed on the Chip Washer (Unchained Labs, US) using the exosome protocol, dried and imaged on the R200 Exoview instrument using Exoview Client. For fluorescent antibodies, the following 3 antibodies were used: CD9-CF488, CD81-CF555 and α-sarcoglycan-CF647 (ORB500418-CF647, Biorbyt). Data was analyzed with Leprechaun Analysis v3.2.1 with the fluorescent cut-offs applied based on particle counts.

microRNAs sequencing of extracellular vesicles from endurance athletes before and after exercise

### EVs isolation for small RNA-sequencing

Small EVs isolated from the serum of 8 endurance athletes before and after exercise were used for small RNA-sequencing (n = 8). Small EVs were isolated using Total Exosomes Isolation Reagent from serum (TEI, #4478360, ThermoFisher). Extraction was performed using 450µl of the serum and mixed with 90µl of the TEI reagent according to the manufacture’s protocol. The pellet with the purified EVs was diluted in 200µl of 1X PBS (pH 7.4, #10010056, ThermoFisher).150µl of the concentrated and purified EVs was submitted for small RNA-seq.

### Sample preparation for small RNA-sequencing

RNA was isolated using the miRNeasy Serum/Plasma Kit (QIAGEN) according to manufacturer’s instructions. QIAseq miRNA Library QC Spike-ins were used for RNA preparation. The library preparation was done using the QIAseq miRNA Library Kit (QIAGEN). A total of 5μl total RNA was converted into miRNA NGS libraries. After adapter ligation, UMIs were introduced in the reverse transcription step. The cDNA was amplified using PCR (22 cycles) and during which indices were added. The samples were then purified. Library preparation was quality controlled using capillary electrophoresis (Tape D1000). Based on quality of the inserts and the concentration measurements, the libraries were pooled in equimolar ratios, quantified using qPCR and then sequenced on a NextSeq 2000 (Illumina Inc.) sequencing instrument according to the manufacturer instructions (single read; read length of 75bp). Raw data was de-multiplexed and FASTQ files for each sample were generated using the bcl2fastq2 software (Illumina inc.). The 19 nt common sequence is: AACTGTAGGCACCATCAAT. The sequencing resulted in 32.6 million reads per sample on average with a median PHRED score of 36 (sequencing accuracy > 99.9%) over the whole read length of all samples (Ewing & Green, 1998).

### Read mapping and differential expression of microRNAs

Raw counts were generated using CLC Genomic Server 21.0.4. Reads were mapped to miRBase version 22 using the “Qiagen miRNA Quantification” workflow with the CLC Genomic Server standard parameters. Raw miRNA count data were filtered to remove lowly expressed miRNAs, retaining only those with a summed count of at least 10 across all samples. Differential expression analysis was conducted using the DESeq2 R package (Love *et al*., 2014), incorporating individual and measurement time in the design matrix. Multiple testing correction was performed using the Benjamini-Hochberg method to control the false discovery rate (FDR). Count data were normalized for downstream visualization using the trimmed mean of M values (TMM) method to account for library size differences and compositional biases (Robinson & Oshlack, 2010).

### Electrical Pulse Stimulation of skeletal muscle cells and tissue

Human skeletal muscle cells (hSKM) were cultured in 12 well plates with Muscle Growth Medium (High-glucose DMEM GlutaMAX with pyruvate (ThermoFisher, Cat. n. 31966) 10% FBS; 1% P/S and 1% L-glutamine). Cells were maintained at 37°C in a humidified incubator with 5% CO_2_. Once cells reached 100% confluency, the medium was replaced with differentiation Myogenic Differentiation Medium (High-glucose DMEM GlutaMAX with pyruvate, 2% horse serum, 1% P/S and 1% L-glutamine) to induce myogenic differentiation. Differentiation was allowed to proceed for one week, with medium changes every 48 hours.

### EPS of human skeletal muscle cells

On the day of EPS treatment and one hour before stimulation, the differentiation medium was replaced with serum-free DMEM to avoid potential interference from serum-derived factors. Electrical pulse stimulation (EPS) was applied using the IonOptix C-Pace system, delivering electrical pulses at 40V, 1 Hz, with a 2-millisecond pulse width for a total duration of 3 hours. For EV collection, the culture medium was immediately collected after EPS. As a control, an additional plate containing the electrodes but without EPS application was subjected to the same media changes and incubation conditions.

### Ex vivo skeletal muscle stimulation

Skeletal muscles were isolated from 8-week-old C57BL/6J mice. The quadriceps were carefully dissected by detaching tendons at the knee and hip joints and removing surrounding connective tissue. Isolated muscles were briefly rinsed in 75% ethanol followed by two washes in phosphate-buffered saline (PBS) to remove debris and potential contaminants. Muscles were placed horizontally between electrodes in EPS plates (IonOptix C-Pace system) and stimulated at 40V, 1 Hz, with a 2-millisecond pulse width for a total duration of 3 hours. Following stimulation, conditioned medium was collected for EV isolation.

### Extracellular vesicle isolation from conditioned media

Conditioned media obtained after EPS was first centrifuged at 2,000 × g for 30 minutes to remove cells and debris. The supernatant was then subjected to EV precipitation using the Total Exosome Isolation Reagent from cell culture media (TEI, # 4478359, ThermoFisher) according to the manufacturer’s instructions. Briefly, TEI reagent was added at a 0.5 volume ratio to the supernatant, mixed thoroughly, and incubated overnight at 4°C. Samples were subsequently centrifuged at 10,000 × g for 60 minutes. The resulting EV pellet was resuspended in phosphate-buffered saline (PBS), typically at a 1:10 dilution relative to the initial conditioned media volume (∼1 ml).

### cAMP treatment of adipocytes and skeletal muscle cells for EV isolation

For cyclic adenosine monophosphate (cAMP) treatment, 500 µM of 8-Br-cAMP (Biolog. Cat n. B 007) diluted in H_2_0 was added to the culture medium (without serum or Penicillin/Streptomycin). A second cAMP treatment was administered two days later to maintain intracellular cAMP levels. EV collection was performed 24 hours after the second cAMP treatment. As a control, an equivalent volume of H_2_O was added instead of cAMP.

### Isolation and culture of murine primary muscle cells

Murine primary muscle cells were isolated from the quadriceps muscles of middle-aged 14 months old Ucp1-Cre X GCAMP5-Td tomato C57BL/6J male mice originally used for a different project where the skeletal muscle tissues were not needed. The mice age correlates to adult humans between 38-47 years old. All animal procedures were planned and performed to minimize pain and suffering, and to reduce the number of animals used in accordance with European, national, and institutional guidelines (European Directive 2010/63/EU; ARRIVE). Isolation and culture of skeletal muscle stem/progenitor cells was based on Soriano-Arroquia et al. (Soriano-Arroquia *et al*., 2017). Briefly, tissues were collected in PBS, washed twice briefly with 70% ethanol, washed twice with PBS and digested in 300-500 µl of pre-warmed enzymatic solution (1.5 U/ml collagenase D, Roche, Cat. n. 11088866001; 2.4 U/ml dispase II, Sigma-Aldrich, Cat. n. D4693; 2.5 mM CaCl_2_, Normapur, Cat. n. VWRC22317.297) diluted in serum-free DMEM/F-12 GlutaMAX media (ThermoFisher, Cat. n. 31331-028). Muscles were mechanically dissociated by cutting the tissue in small pieces with dissecting scissors and incubated in a water bath at 37°C to allow enzymatic digestion for 1 hour, tapping the tissue every 10 minutes. After digestion, cells were re-suspended in 10 ml of seeding media and filtered using a 100 µm cell strainer. Next, cells were centrifugated at 443 x g for 5 minutes at room temperature, supernatant discarded, re-suspended in seeding media for expansion or further experiments.

After expansion, murine primary muscle stem/progenitor cells were grown in Muscle Growth Media (High-glucose DMEM GlutaMAX with pyruvate (ThermoFisher, Cat. n. 31966) 10% FBS; 1% P/S and 1% L-glutamine). When confluent, cells were allowed to differentiate into myotubes or muscle-derived adipocytes. For myogenic differentiation, growth media was change to Myogenic Differentiation Media (High-glucose DMEM GlutaMAX with pyruvate, 2% horse serum, 1% P/S and 1% L-glutamine) and cultured seven days in a humidified incubator at 37°C and 5% CO_2_ until myotubes were visible under the microscope. To allowed brown/beige adipogenic differentiation of murine primary muscle stem/progenitor cells, media was changed to BA Differentiation Media (High-glucose DMEM GlutaMAX, ThermoFisher, Cat. n. 61965; 10% FBS, 1% P/S, 20 nM insulin and 1 nM triiodothyronine) for two days. Next, media was changed to BA Induction Media (BA differentiation media supplemented with 0.5 mM isobutylmethylxanthine (IBMX) and 1 µM dexamethasone) for two days and finally changed again to BA Differentiation Media for five more days until mature adipocytes were visible under the microscope.

### In vitro overexpression of microRNAs

For overexpression of miR-196a-5p and miR-342-3p, lentiviral particles encoding the corresponding human precursor microRNA sequences were used. A non-targeting scrambled microRNA sequence served as a negative control. The miRNA expression cassettes were cloned into a lentiviral backbone under the control of the CMV promoter. Cells were transduced at approximately 70-80% confluency in growth medium containing lentiviral particles. After 24 hours the virus-containing medium was replaced with fresh growth medium and cells were allowed to recover before induction of adipogenic or myogenic differentiation. Overexpression of the respective mature microRNAs was confirmed by RT-qPCR using TaqMan Advanced miRNA assays.

**Table 2:**
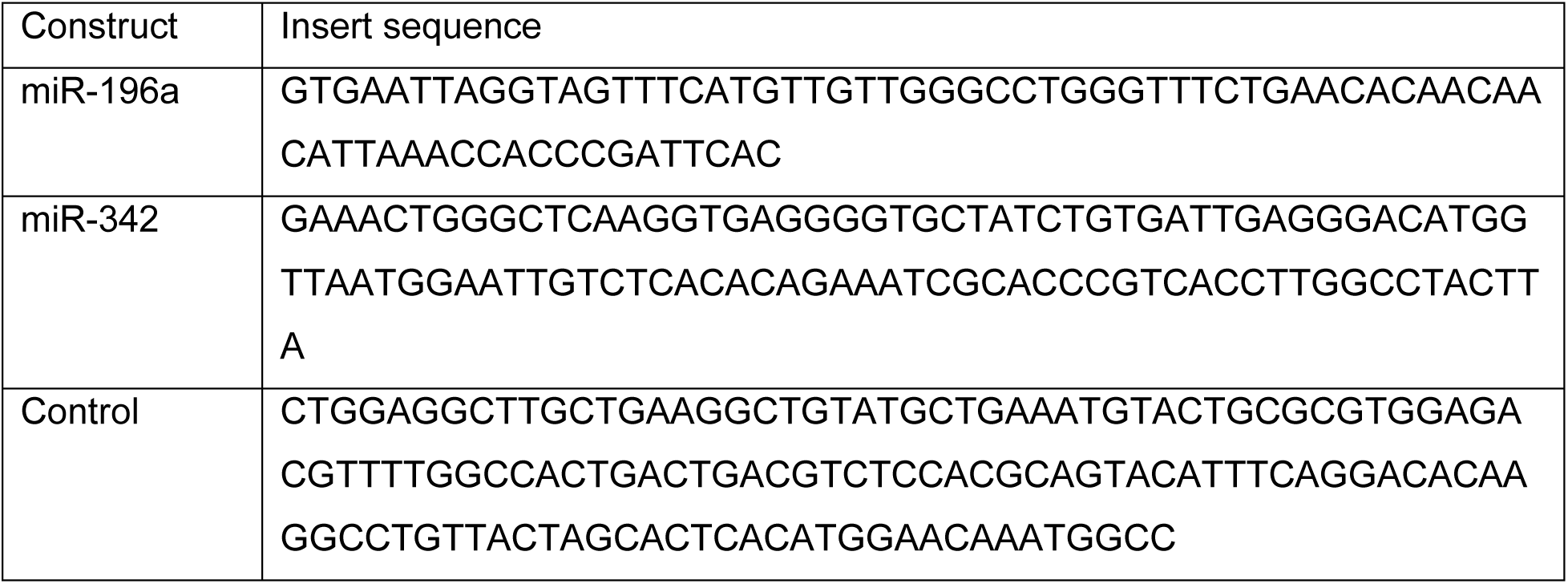
Insert sequences for the constructs used for lentiviral transduction.

### RNA isolation and quantitative real-time PCR

RNA isolation of cells lysate or EVs isolated from cells conditioned media was performed using Trizol-Chloroform standard protocol and quantified by Nanodrop spectrophotometer.

For the cell lysate, cDNA was synthesize using ProtoScript II First Strand cDNA Synthesis Kit (New England Biolabs, Cat. n. E6560). Quantitative PCR was performed using Luna Universal qPCR Master Mix (New England Biolabs, Cat. n. M3003) using a QuantStudio Real-Time PCR System (ThermoFisher). Ct values were normalized to the housekeeping gene (Hprt, beta-actin or TBP as stated in each figure) and values shown as ΔCt. The primer’s sequences used in this study are included in suppl. table 2. For qPCR quantification and validation of *UCP1* mRNA levels in hWAs after serum and EV treatment, TaqMan assays and TaqMan Fast Advanced Master Mix with UNG (4444963, ThermoFisher) were used according to the manufacture’s protocols.

For cells-derived EVs, microRNA quantification was performed using TaqMan Advanced miRNA cDNA Synthesis Kit (A28007, ThermoFisher) and TaqMan Fast Advanced Master Mix with UNG (4444963, ThermoFisher) according to the manufacture’s protocols. miRNA Ct values were normalized to an internal control (miR-320a-3p). We selected miR-320a-3p as the best endogenous microRNA suggested by ThermoFisher with the lowest variability between treatment in our samples. TaqMan primers references are shown in suppl. table 3.

### Proteomics of human serum-derived EVs before and after exercise

#### Sample preparation for proteomics analysis

For proteomics analysis, 40 µl of serum-isolated EVs were lysed by adding 40 µl urea lysis buffer (4 M urea, 10 mM Bond-Breaker TCEP solution [Thermo], 30 mM CAA [Merck] in Tris-HCl pH 8.0) followed by incubating the samples on a thermo shaker at 800 rpm and room temperature for 30 min. A modified version of the single-pot solid-phase-enhanced sample preparation (SP3) was used for sample cleanup and trypsin digestion (https://doi.org/10.1038/s41596-018-0082-x). In brief, 2 µl of ready-prepared SP3 magnetic beads (SpeedBeads magnetic carboxylate modified particles [Merck]) and 100 µl of acetonitrile (ACN, [VWR]) were added to each sample, followed by incubation on a thermo shaker at 1,000 rpm and room temperature for 5 min. Samples were then incubated for 2 min on a magnetic rack, and the supernatant was removed. Then, samples were washed 3 times with 180 µl of 80 % ethanol and air-dried. Magnetic beads were dissolved in 10 µl digestion buffer (0.1 µg trypsin/Lys-C [Promega] in 50 mM TEAB) and digested at 37 °C and 800 rpm in a thermo shaker for 18 hours. On the next day, 200 µl of pure ACN was added, and samples were again incubated at 800 rpm and room temperature for 5 minutes. Samples were transferred onto a magnet, the supernatant was discarded, and beads were rinsed with 200 µl ACN. Peptides were eluted from beads with 10 µl of 5 % ACN in LC-MS-grade water [VWR] and transferred to fresh tubes. Samples were acidified by adding 2 µl of 10 % formic acid [Sigma], and peptide concentrations were determined using a Nanodrop One system [Thermo]. Finally, 200 ng of peptides were loaded onto an Evotip Pure [Evosep] trap column following the manufacturer’s protocol.

### LC-MS/MS analysis

Proteomics samples were measured with a liquid chromatography-tandem mass spectrometry system consisting of an Evosep One chromatographic system [Evosep] and a timsTOF Pro 2 mass spectrometer (Bruker Daltonics). Peptides were separated by the preinstalled 40-samples-per-day whisper method using 31 min chromatographic gradients. Peptides were separated by an Aurora Elite CSI analytical column [Ionopticks] of 15 cm length and 75 µm inner diameter and filled with 1.7 µm C18 particles. The LC system was online-coupled to the mass spectrometer. Eluting peptides were transferred directly into the MS via a Captive Spray nano-electrospray ion source operated at a constant spray voltage of 1.5 kV. Samples were measured in data-independent acquisition parallel accumulation-serial fragmentation (diaPASEF) mode. The dual TIMS analyser operated with a 100% duty cycle using 100 ms accumulation and ramping times, respectively. Peptides were isolated from a mass-to-charge range of 400 – 1000 m/z and an ion mobility range of 0.70 – 1.35 Vs/cm^2^. The isolation space was separated into 24 equally sized PASEF windows of 25 m/z each. The collisional energies were stepwise increased from 24 eV at 0.7 Vs/cm^2^ to 49 eV at 1.35 Vs/cm^2^.

### Data processing and statistical analysis of proteomics data

Proteomics data of exercise-induced EVs were processed by DIA-NN (version 1.8.1, (Demichev *et al*., 2020). The library-free search strategy was utilised to predict a spectral library and was used for peptide identification and quantification. The SWISS-PROT Homo sapiens FASTA database downloaded from UniProt (version from 2023-02-15) was used to generate the spectral library *in-silico*. Cysteine carbamidomethylation was set as a fixed modification, and trypsin was selected as the digestion enzyme. Maximum one missed cleavage was allowed. Mass accuracies were determined beforehand and were fixed to 1.3e-05 (MS2) and 1.1e-05 (MS1), respectively. Precursor peptides were filtered at an FDR<1%. Protein group intensities were normalised by the MaxLFQ algorithm (Cox *et al*., 2014), considering proteotypic peptides only for quantification.

The statistical analysis was performed with the Perseus software suite (v. 1.6.15, Tyanova et al. 2016). In both data sets, protein LFQ intensities were Log_2_ transformed and replicates with reduced identifications removed from the analysis. Protein groups with less than 70 % data completeness in at least one condition were removed. Missing values were replaced sample-wise by random value drawings from 1.8 standard deviations downshifted and 0.3 standard deviations broad normal distribution.

### Statistics

Two-tailed unpaired or paired t-tests were used for statistical differences between two groups. For multiple comparison, either one-way ANOVA and Tukey’s multiple comparisons test, or Two-way ANOVA and Sidak’s multiple comparisons test, with 95% confidence interval, were used unless otherwise stated. p-value < 0.05 was considered as statistically significant. Error bars are shown as the Standard Error of the Mean (S.E.M.). Exact details of the specific statistics are shown in each figure legend.

## Results

### Post-exercise serum from endurance athletes increases *UCP1* transcription in human white adipocytes

To screen for candidate exercise-induced microRNAs that mediate skeletal muscle and thermogenic adipose tissue crosstalk in humans, and to investigate whether the training background influences this crosstalk, we isolated the serum of sedentary individuals, endurance athletes, natural bodybuilders and sprinters before and after a bout of graded cycle ergometer to exhaustion (Schranner *et al*., 2021). Direct treatment of mature hWAs with the same volume of post-exercise serum isolated from endurance athletes, but not from sedentary individuals, bodybuilders and sprinters, resulted in a 2.8-fold increase of *UCP1* mRNA levels (Fig. 1A). We hypothesized that the endurance-trained phenotype may be associated with exercise-induced EV-miRNA changes that stimulate *UCP1* transcription. To test this, we isolated serum small EVs from each group (Figs. 1B-D) and treated the cells with the same amount of EVs (Fig. 1E). We did not find significant differences between pre-versus post-exercise serum-EV treatment across the four groups. However, treatment with the post-exercise EVs from endurance athletes significantly increased the *UCP1* mRNA levels in hWAs in comparison to the post-exercise treatment in the other groups (Fig. 1E). In order to further investigate whether exercise-induced muscle-specific EVs promote *UCP1* expression in hWAs, we used an in vitro model of acute exercise based on Pillon et al. (Pillon *et al*., 2020). The model involved EPS of human skeletal myotubes at day 7 post-differentiation (Fig. 1F). Immediately after EPS, muscle-EVs were isolated from the conditioned media for adipocytes treatment. Treatment of hWAs with muscle-derived EVs without any treatment resulted in a significant 8.5-fold increase in *UCP1* levels (p = 0.01), and 14-fold increase in *UCP1* levels using muscle-derived EVs after EPS (p = 0.0003) (Fig. 1G). Muscle-EVs collected after EPS did also enhance the expression of other thermogenic markers such as *PGC1α* (1.3-fold, p = 0.03) and *DIO2* (2.4-fold, p = 0.04) when compared to EVs control, and non-significant differences in *PPAR*γ and *CIDEA* (Fig. 1H-J). Additional EV characterization revealed a small population of serum-derived CD63+ EVs positive for alpha sarcoglycan (SGCA), a potential marker of muscle-derived EVs (Guescini *et al*., 2015), highlighting the presence of muscle EVs in the human serum (Fig. 1L). Overall, these results indicate that muscle EVs are able to enhance thermogenic markers, especially *UCP1,* in hWAs, which effect is boosted under exercise-mimicking conditions.

**Figure 1.**
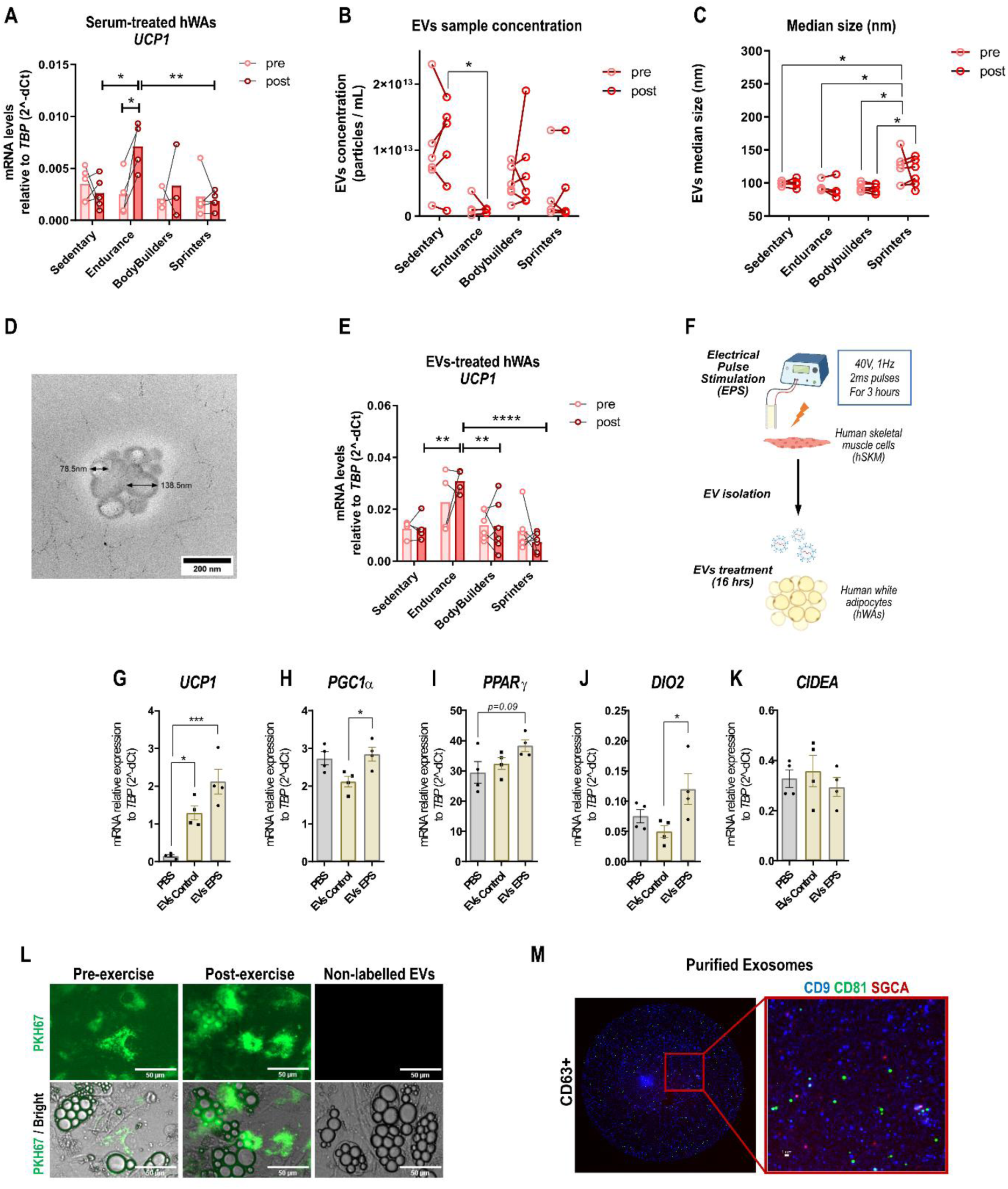
*UCP1* mRNA levels in human white adipocytes (hWAs) treated with serum (A) from sedentary controls, endurance athletes, bodybuilders and sprinter athletes. Concentration (B) and median size (C) of serum-derived small extracellular vesicles using Nanoparticle Tracking Analysis. Electron microscopy imaging of serum-derived small EVs from post-exercise serum of endurance athletes (D). *UCP1* mRNA levels in hWAs treated with serum small EVs (E). Two-way ANOVA, Sidak’s multiple comparisons test, n = 3-6. Schematic of electrical pulse stimulation protocol and EV transfer to hWAs (F). mRNA expression of the thermogenic markers *UCP1* (G) *PGC1α* (H), *PPARγ* (I) DIO2 (J) and CIDEA (K) in hWAs treated with either non-stimulated muscle EVs (EVs Control) or muscle EVs collected after EPS (EVs EPS). PBS treatment was used as a negative control. Forskolin treatment was performed in parallel as a positive control for *UCP1* expression (data not shown). One-way ANOVA, Tukey’s multiple comparisons test, n = 4. PKH67 staining of serum-derived EVs up-taken by treated hWAs (L). ExoView analysis showing immuno-captured EVs positives for CD63 co-expressing CD9 (blue), CD81 (green) and α-sarcoglycan (SGCA, red) (L). Error bars show SEM. * - p < 0.05; ** - p < 0.01; *** - p < 0.005.

### Small RNA-sequencing reveals differentially expressed microRNAs in endurance athletes after exercise

We next focused further investigations to screen for exercise-induced muscle-specific microRNAs that contribute to the increased expression of *UCP1* in hWAs. To identify candidate microRNAs that play a role in skeletal muscle and adipose tissue crosstalk, small RNA-sequencing was performed on purified serum EVs from 8 endurance athletes before and after exercise (Fig. 2A). From a total of 2632 microRNAs, 1130 microRNAs were detectable in at least one sample. Using a raw count threshold > 10, absolute Log_2_ fold-change threshold > 0.2 and an FDR adjusted p-value < 0.05, we identified 16 microRNAs significantly upregulated and 10 microRNAs significantly downregulated after exercise (Fig. 2B and suppl. table 1). In order to narrow down potential candidates that represent crosstalk between skeletal muscle and thermogenic adipose tissue, we performed a systematic analysis using PRISMA guidelines (Page *et al*., 2021) of published studies investigating microRNAs in skeletal muscle, adipose tissue, serum or plasma under certain conditions or stimuli such as exercise, cAMP treatment and cold exposure. The database generated from the systematic collection of previous omics data was manually curated and it is fully available to the scientific community in the supplementary information (suppl. Fig. 1A). Using our systematic database, we filtered those microRNAs and EV-derived microRNAs enriched at baseline levels from skeletal muscle cells or brown/beige adipocytes, or upregulated by either exercise, cold exposure or cAMP stimulation (suppl. Fig. 1B). We next checked which of those microRNAs collected from the database are also upregulated in the post-exercise serum of endurance athletes according to our small RNA-sequencing. From muscle derived microRNAs, only miR-143-3p (log_2_ FC = 0.42, padj = 0.02) and miR-196b-5p (log_2_ FC = 0.47, padj = 0.03) were significantly upregulated in the EVs from endurance athletes after exercise (Fig. 2C, 2E). Particularly, miR-143-3p was up-regulated in human muscle-derived fibroblasts EVs from patients with Duchenne muscular dystrophy (DMD) (Zanotti *et al*., 2018) and in circulating EV-derived microRNAs after acute exercise in healthy men (Nielsen *et al*., 2014). In addition, miR-196b-5p was enriched in C2C12 murine myotubes and derived EVs (Garcia-Martin *et al*., 2022), in the vastus lateralis muscle from women with polymyositis and dermatomyositis after endurance training (Boehler *et al*., 2017), and in older males (66.6 ± 1.1 years old) after acute resistance exercise (Zacharewicz *et al*., 2014).

**Figure 2.**
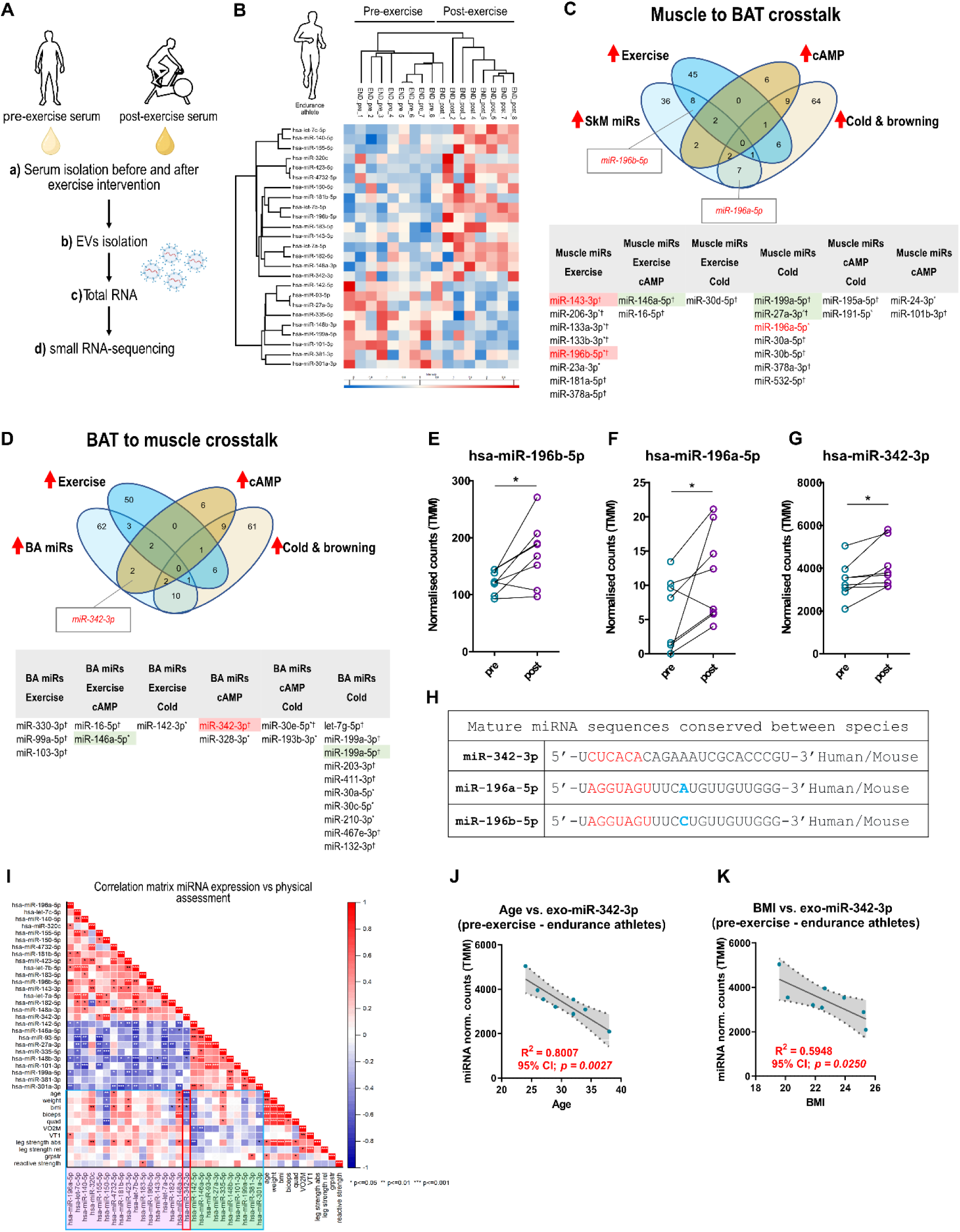
Schematics used for the isolation of serum-derived small EVs from endurance athletes before and after exercise (A). Heatmap showing differentially expressed microRNAs (B). Venn diagrams showing microRNAs (*) and microRNAs contained in EVs (^†^) enriched in skeletal muscle cells (C) or brown adipocytes (D), that are upregulated in the serum/plasma or cell conditioned media after exercise, cAMP or cold stimuli according to previous published studies. Highlighted in red (up-regulated) or in green (down-regulated) are shown those microRNAs differentially expressed in serum-derived small EVs from endurance athletes after exercise based on our small RNA-sequencing. TMM normalized count levels of candidate microRNAs before and after exercise (E-G). Two-tailed paired t-test. n = 8. Error bars show SEM. * - p < 0.05; ** - p < 0.01; *** - p < 0.005. The three candidate microRNAs are evolutionary conserved between human and mice. miR-196a-5p and miR-196b-5p share the same seed sequence (shown in red), which indicates similar mechanistic effects through the repression of the common target genes (H). Correlation matrix of differentially expressed microRNAs and physical parameters of endurance athletes (I). Matrix was performed using OriginPro, Version 2023 (OriginLab Corporation, Northampton, MA, USA). Pearson’s correlation of miR-342-3p normalized counts versus age (J) and BMI (K). n = 8.

Interestingly, miR-196a-5p, a cold-induced microRNA required for browning of subcutaneous white adipocytes (Mori *et al*., 2012a), was also enriched in C2C12 myotubes based on Garcia-Martin et al. omics data (Garcia-Martin *et al*., 2022). Both miR-196b-5p and miR-196a-5p share the same seed sequence (Fig. 2H) and are predicted to target the Hox gene *HOXC8*, whose inhibition induces browning of white adipocytes by de-repressing C/EBPβ (Mori *et al*., 2012a). Hence, we checked the expression of miR-196a-5p in our small RNA-sequencing data. We found that miR-196a-5p was significantly higher expressed after exercise (log_2_ FC = 1.25, non-adjusted paired p-value = 0.044) (Fig. 2F). Although miR-196a-5p was not retained in the final DESeq2-filtered list of differentially expressed microRNAs due to the low mean count abundance, it was examined as a hypothesis-driven candidate based on its close relationship to miR-196b-5p and its established role in adipocyte browning. A recent study shows that BAT releases EVs during exercise with a cardioprotective effect, emerging BAT as an alternative source of EVs after exercise (Zhao *et al*., 2022). Therefore, we also screened for BAT-derived microRNAs to investigate potential BAT-to-muscle interactions. From the list of microRNAs derived from brown adipocytes, only miR-342-3p overlapped with those up-regulated in the serum of endurance athletes after exercise (Fig. 2D, 2G). According to our database, miR-342-3p was enriched in mouse immortalized brown adipocytes (Garcia-Martin *et al*., 2022) and in EVs released from cAMP-treated murine brown adipocytes versus non-treated brown adipocytes (Chen *et al*., 2016). Notably, miR-342-3p expression negatively correlated with both age and BMI (Fig. 2I-K). We therefore focused further studies on miR-196a-5p, along with miR-196b-5p and miR-342-3p. None of these microRNAs were previously investigated for their role in skeletal muscle and adipose tissue crosstalk during exercise through small extracellular vesicles.

### In vitro model of exercise promotes the release of muscle-specific EVs enriched in miR-196a-5p

To investigate whether muscle cells release small EVs enriched in our candidate microRNAs under exercise mimicking conditions, we next checked the expression of these microRNAs in muscle-derived small EVs after EPS versus non-stimulated control EVs. Stimulation with EPS showed a 2.5-fold significant increase of miR-196a-5p expression in human myotubes-derived EVs (p = 0.03) (Fig. 3A). miR-196b-5p showed non-significant differences (7.5-fold, p = 0.1) (Fig. 3B). We did not find relevant differences in the expression of miR-342-3p after EPS in vitro (Fig. 3C). Similarly, ex vivo analysis of EVs released by murine quadriceps after EPS showed significant increase of miR-196a-5p (1.7-fold, p = 0.007), but no changes in miR-196b-5p expression (p = 0.90), while miR-342-3p was significantly increased (1.6-fold, p = 0.03) (Fig. 3D-F) when compared to the non-stimulated control EVs. Since there was an increase of EV-derived microRNA-342-3p ex vivo but not in vitro, we hypothesized that miR-342-3p might be specific from other muscle cell types such as fibro/adipogenic progenitor (FAPs)-derived adipocytes. To corroborate the muscle cell type specificity of our candidate microRNAs, a mixed population of murine primary stem/progenitor muscle cells was isolated from the quadriceps of mice and differentiated under either myogenic or adipogenic media (Suppl. Fig. 2A). Only satellite cells will differentiate into myotubes under myogenic conditions, while only FAPs will differentiated into adipocytes under adipogenic conditions (Joe *et al*., 2010; Uezumi *et al*., 2011). Differentiated myotubes or mature FAPs-derived adipocytes were treated with 0.5mM cAMP at day 7 and day 9 post-differentiation with a resting day between treatments (Suppl. Fig. 2A). cAMP treatment of myotubes, but not muscle-derived adipocytes, released EVs enriched in miR-196a-5p (5-6-fold, p = 0.0001) and miR-196b-5p (4.1-fold, p = 0.04) (Suppl. Fig. 2B, 2C). Compared to miR-196a-5p, the basal levels of miR-196b-5p in myotubes-derived EVs was lower (Suppl. Fig. 2C). Interestingly, miR-342-3p levels decreased after cAMP treatment in myotubes-derived EVs (0.3-fold, p = 0.0007) (Suppl. Fig. 2D), suggesting a different cell specificity and a preference of muscle cells to accumulate or uptake this microRNA under cAMP stimuli.

**Figure 3.**
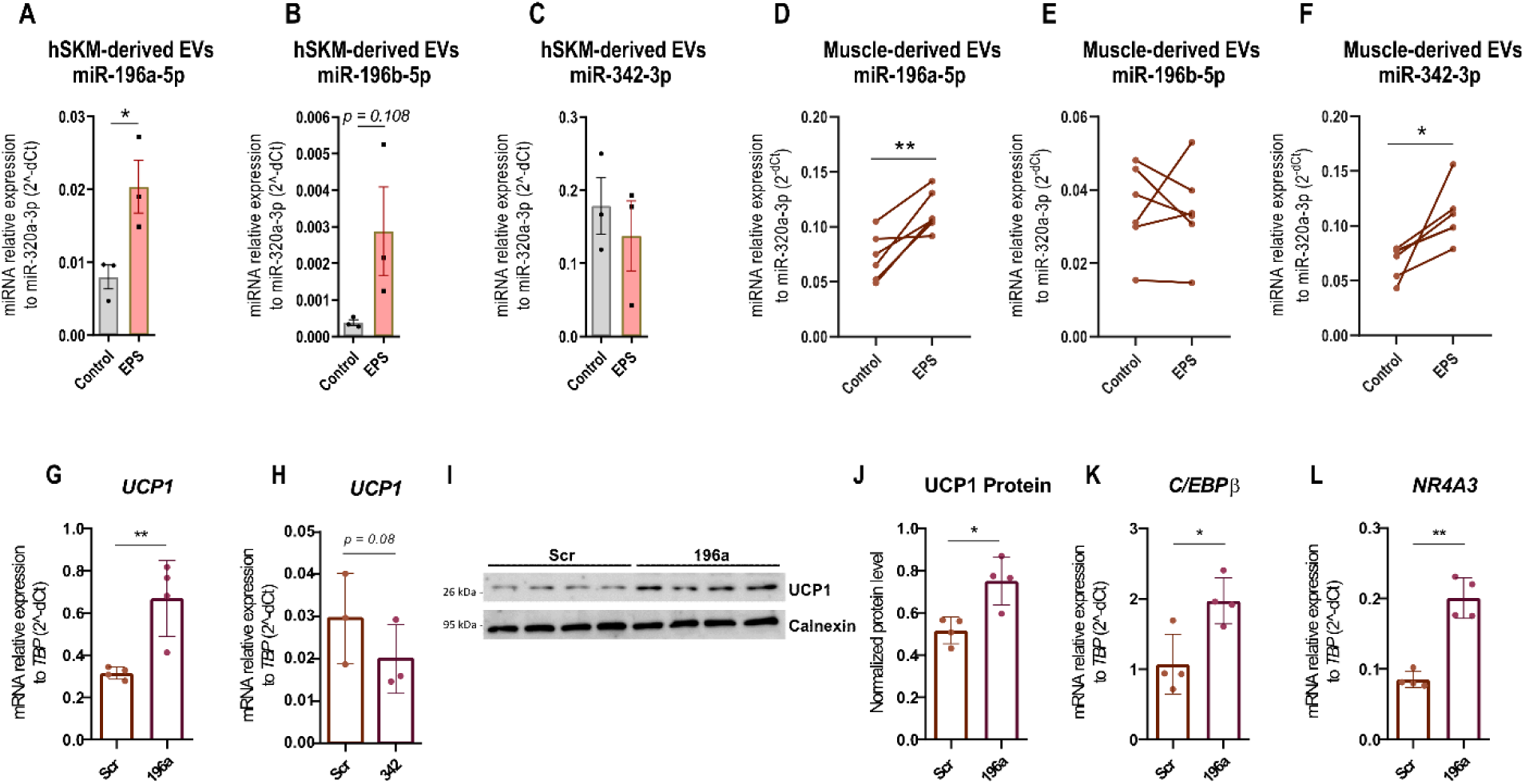
Human skeletal muscle cells release EVs enriched in miR-196a-5p (A) but not in miR-196b-5p (B) or miR-342-3p (C) after EPS. Two-tailed unpaired t-test compared to the mock control, n = 3. Ex vivo microRNAs expression in EVs isolated from murine quadriceps after EPS compared to non-stimulated tissues (Control) (D-F). Two-tailed paired t-test compared to the mock control, n = 5. *UCP1* mRNA levels after miR-196a (G) or miR-342 (H) lentivirus overexpression in hWAs. Western blot (I) and protein quantification (J) showing UCP1 protein abundance in hWAs after miR-196a overexpression. *C/EBPβ* (K) and *NR4A3* (L) mRNA levels in hWAs after miR-196a overexpression compared withscrambled control (Scr). Two-tailed unpaired t-test compared to Scrambled control (Scr), n = 4. Error bars show SEM. * - p < 0.05; ** - p < 0.01; *** - p < 0.005.

To find out whether miR-342-3p directly regulates *UCP1* transcription, we overexpressed miR-342-3p, along with miR-196a-5p as a positive control, in hWAs (Suppl. Fig. 2E-G). Our results showed an increased expression of *UCP1* at the mRNA and protein levels by miR-196a-5p lentivirus overexpression in mature hWAs (Fig. 3G, 3I, 3J) as previously shown by Mori et al. (Mori *et al*., 2012a). However, *UCP1* was not significantly increased after miR-342-3p overexpression (p = 0.08) (Fig. 3H), indicating that among all the microRNAs upregulated after exercise, microRNA-196a-5p is the strongest candidate to induce *UCP1* expression in hWAs. Consistent with the previously described miR-196a–HOXC8–C/EBPβ pathway (Mori *et al*., 2012a), miR-196a-5p overexpression reduced *HOXC8* expression (Suppl. Fig 2H) and increased *C/EBPβ* (2-fold, p = 0.02) (Fig. 3K), supporting activation of a thermogenic transcriptional program. In addition, miR-196a-5p increased *NR4A3* expression (2.42-fold, p = 0.0003) (Fig. 3L), an exercise-responsive nuclear receptor (Smith *et al*., 2025), suggesting that miR-196a-5p may not only induce classical browning markers but also promote a broader exercise-associated response in hWAs.

### Multi-omics correlation and enrichment analysis of upregulated microRNAs and proteomics in serum-derived EVs from endurance athletes after exercise

EVs contain intracellular and membrane proteins that can be used as signatures or footprints of their cellular origin (Chen *et al*., 2022). Based on this concept, we performed proteomics analysis to screen for tissue-specific markers and explore the potential source of microRNAs upregulated after exercise, including our candidate microRNAs. For that, we isolated EVs from the same participants for proteomics analysis (pre versus post exercise, n=6 per group). EV proteomics identified a total of 1395 proteins across all samples. Samples with less than 650 identified proteins were considered outliers and excluded from the analysis along with their respective paired samples. Outlier removal resulted in the exclusion of 7 paired samples, making a total of n=4 sedentary controls; n=3 bodybuilders; n=5 endurance athletes; and n=5 sprinters for statistical and enrichment analyses. Complete information about significantly regulated proteins by time and by training group can be found in the supplementary information (suppl. data 3).

To characterize possible cellular sources of the microRNAs induced after exercise, we performed an integration analysis of the small RNA-sequencing and proteomics data. First, the normalized counts of the up-regulated microRNAs in endurance athletes after exercise were correlated with the corresponding proteins from each participant. A total of 257 out of 1395 proteins (18.4%) positively correlated with the expression of at least one microRNA (Pearson’s correlation, p ≤ 0.05, suppl. data 4). The list of positively correlated proteins was used for functional enrichment analyses using FunRich software (Pathan *et al*., 2015). 93% (239 proteins) overlapped with the Vesiclepedia database (Chitti *et al*., 2024) (Fig. 4A). From those 239 proteins, 62.4% were previously identified in exosomes (Fig. 4B), supporting that most of the proteins identified in our proteomics data have been previously detected in EVs from other studies and suggesting the reproducibility of our proteomics method. Since we were specifically interested in the identification of EVs mediating skeletal muscle and adipose tissues crosstalk, we particularly checked for those proteins present in both tissues. 68% of the 239 proteins were previously identified in skeletal muscle EVs, 22.4% in pre-adipocytes EVs and only 2.9% in adipose tissue EVs (Fig. 4C). As already reported by other studies (Ismaeel *et al*., 2023), we could not detect the SGCA protein in serum-derived EVs, presumably due to overrepresentation of abundant proteins masking lower abundant ones, emphasizing the need for identifying alternative muscle-EV-specific markers. Intriguingly, the fatty acid binding protein 4 (FABP4 or aP2) was present in the three groups (Fig 4D). FABP4 is a cytoplasmatic protein highly enriched in mature adipocytes that regulates fatty acid transport and uptake (Storch & McDermott, 2009), but also present to a lesser extend in endothelial cells, smooth muscle cells and macrophages (proteinatlas.org). Therefore, we used FABP4 as potential adipocytes-derived EV marker for further exploration.

**Figure 4.**
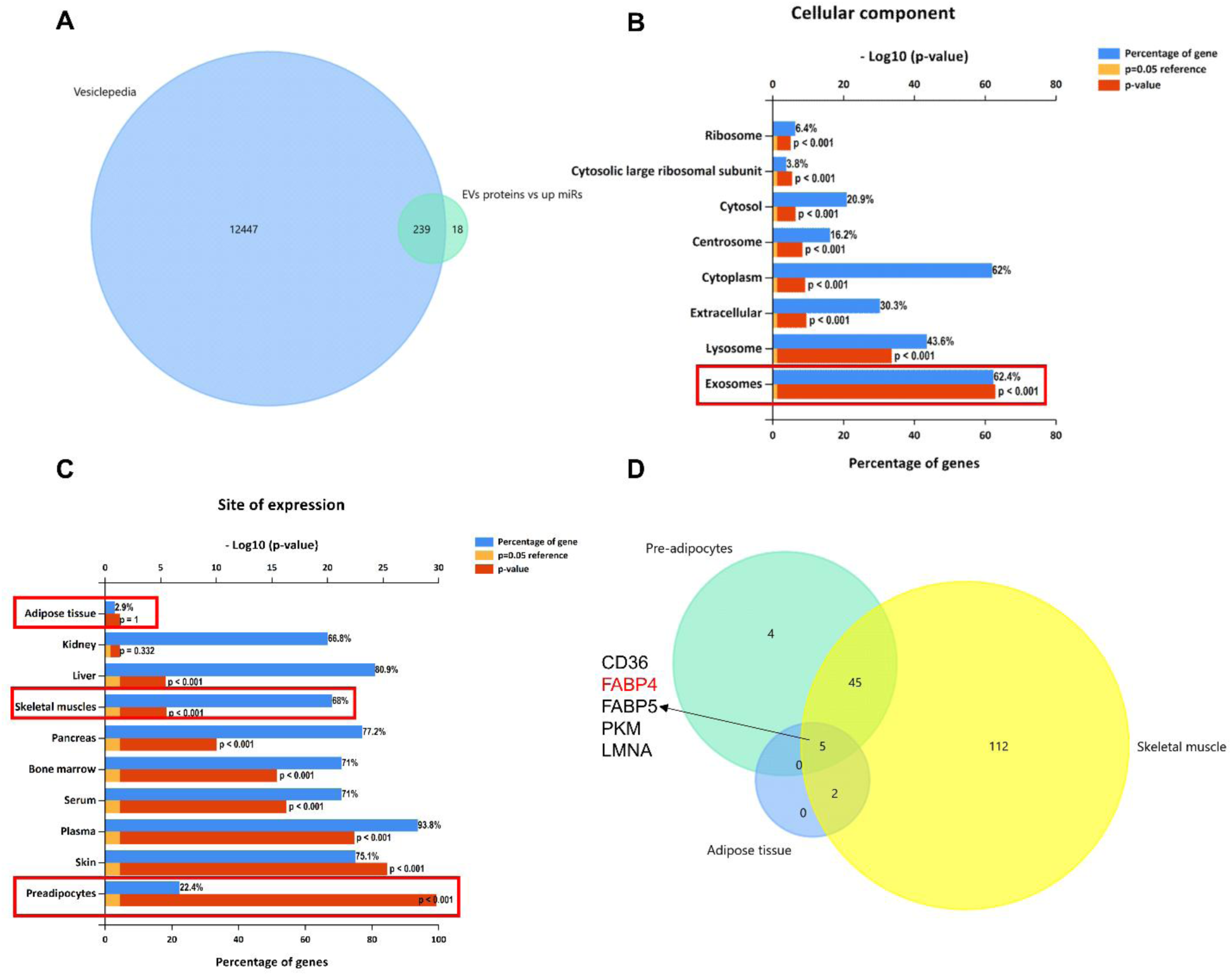
Functional Enrichment Analysis using FunRich software of small RNA-sequencing-proteomics integration analysis. From a total of 257 EVs proteins that positively correlated with the expression of upregulated microRNAs after exercise in endurance athletes, 93% (239 proteins) overlapped with the Vesiclepedia database (A). Cellular component enrichment analysis showing a 62.4% overlap with proteins previously found in exosomes (B). Site of expression analysis of the most representative organs and tissues (C). Venn diagram showing the overlap of EV-proteins previously found in skeletal muscle, pre-adipocytes and adipose tissue (D).

### Human white and brown adipocytes release EVs enriched in miR-342-3p with myogenic potential

Based on our multi-omics analysis, we next investigated whether FABP4+ EVs could be released from adipocytes containing our candidate microRNAs. We hypothesised that exercise may induce the release of FABP4+ EVs from stimulated adipocytes which are enriched in microRNAs with myogenic potential. Our proteomics analysis showed that FABP4 is particularly increased in serum-derived EVs from endurance athletes after exercise, but not in sedentary individuals, bodybuilders or sprinters (Fig. 5A). We next analysed the list of up-regulated microRNAs that positively correlated with FABP4 protein levels. We found that only miR-342-3p positively correlated with FABP4 protein levels in EVs from endurance athletes (R^2^ = 0.6376, p-value = 0.0056) (Fig. 5B) but not significantly correlated with miR-196a-5p (R^2^ = 0.2245, p-value = 0.1665) or miR-196b-5p (R^2^ = 0.1711, p-value = 0.2347) (suppl. Fig. 5A, B). We therefore investigated whether white adipocytes release EVs enriched in miR-342-3p. Of note, cAMP stimulation of hWAs resulted in 2.4-fold increased expression of miR-342-3p (p = 0.02) in the released EVs compared to non-stimulated control cells (Fig. 5D). Similarly, miR-342-3p also showed a strong trend in EVs derived from human brown adipocytes (7.01-fold, p =0.055) (Fig. 5E). We did not find a significant change in the expression of miR-196a-5p in white or brown adipocyte-derived EVs (Suppl. Fig. 5C, 5D). Furthermore, microRNA-196b-5p was not detectable by qPCR in either the control nor cAMP-stimulated adipocytes. These results suggest that white and brown adipocytes could be a source of microRNA-342-3p upon cAMP stimulation. Since we aimed to find out bi-directional crosstalk between skeletal muscle and adipose tissue, we next investigated whether miR-342-3p regulates myogenic markers in vitro. To investigate this, hSKMs were transduced with miR-342 lentivirus. Cells were collected at Day 2, 5 and 7 during myogenic differentiation and myogenic markers were detected by RT-qPCR. Overexpression of miR-342 resulted in increased levels of PRDM16 (1.51-fold, p = 0.03) and *MYOD* at day 2 (4.79-fold, p = 0.056) (Fig. 5F) and increased levels of PGC1α (1.5-fold, p = 0.04) *MYOG* (which encodes for Myogenin) at day 7 (2.25-fold, p = 0.04) (Fig. 5G). Together, these findings suggest that cAMP-stimulated white and brown adipocytes can release EVs enriched in miR-342-3p, and that miR-342-3p overexpression is associated with increased expression of mitochondrial biogenesis and myogenic markers during hSKMs differentiation.

**Figure 5.**
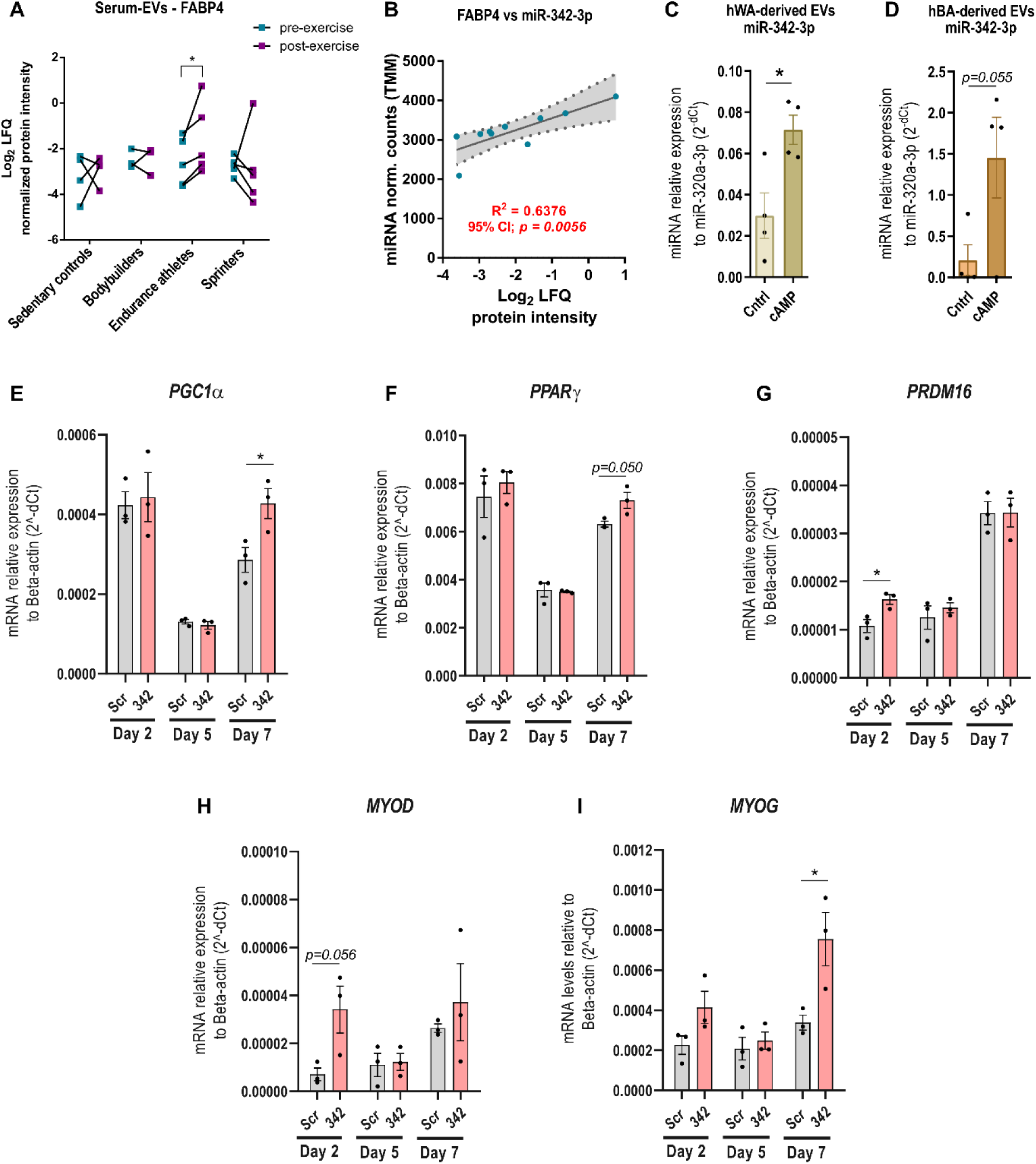
Proteomics analysis showing increased levels of FABP4+ after exercise in EVs isolated from endurance athletes but not from sedentary controls, bodybuilders or sprinters (A). Two-way ANOVA, Sidak’s multiple comparisons test. n = 3-5. Pearson’s correlation of miR-342-3p normalized counts versus FABP4 Log_2_ normalized protein intensity (B). n = 8. Real-Time qPCR expression of miR-342-3p in EVs released from hWAs (D) or hBAs (E) after cAMP treatment compared to non-treatment control (Ctrl). Two-tailed paired t-test. n = 4. *PGC1α* (E), *PPARγ* (F), *PRDM16* (G), *MYOD* (H) and *MYOG* (I) mRNA expression of hSKM transduced with miR-342 at days 2, 5, and 7 post myogenic differentiation compared to scrambled control (Scr). Two-tailed unpaired t-test, miR-342 versus Scr. n = 4. Error bars show SEM. * - p < 0.05; ** - p < 0.01; *** - p < 0.005.

## Discussion

Induced thermogenesis of subcutaneous white adipose tissue (scWAT) has been proposed as a potent therapeutic target against metabolic diseases such as obesity, especially in individuals with low or non-functional brown adipose tissue (BAT). While there is strong evidence showing that exercise induces WAT browning in rodents, there is still no consensus about its effect in humans. To date, the human research shows contradictory findings to this respect: Otero-Díaz et al. showed an increased expression of UCP1 and other thermogenic markers in scWAT after 12-week of exercise training in non-diabetic sedentary individuals (Otero-Díaz *et al*., 2018); other studies have shown no conclusive changes of thermogenic markers in scWAT after long-term training (Norheim *et al*., 2014; Tsiloulis *et al*., 2018). Similarly, a recent study published by us show no effect in cold-induced thermogenesis after 6-week of training in sedentary individuals (Braunsperger *et al*., 2026). Interestingly, some rodent studies show antagonistic inhibition of BAT but increased thermogenesis in WAT after chronic exercise in obese mice (Wu *et al*., 2014), suggesting a more relevant role of WAT thermogenesis during exercise.

The fact that most of the exercise studies in rodents have been performed below thermoneutral conditions (where thermogenic adipose tissue is constantly activated) and that human interventions are very limited and have been mostly performed in healthy or obese sedentary individuals, raises the question of whether other training backgrounds and environmental stimuli might result in different outcomes in humans. There is only one study in humans comparing training backgrounds. Vosselman et al. found lower BAT glucose uptake after cold exposure in endurance athletes compared to sedentary controls, and no differences of WAT browning at thermoneutrality between both groups (Vosselman *et al*., 2015). Still, since only glucose uptake is measured by ^18^F-FDG PET/CT scans, it remains unanswered whether thermogenic adipose tissue from endurance athletes might have a higher preference of fatty acids or other substrates during cold-induced thermogenesis. Likewise, the effects of acute exercise to exhaustion in adipose tissue thermogenic on markers has not been tested yet. Interestingly, it has been previously shown that just a bout of aerobic incremental exercise until achieving the “anaerobic” threshold (∼80-85% of VO_2_max) induces a greater release of circulating myokines and adipokines such as FGF-21 and follistatin, known to induce browning, compared to lower intensities (∼50-60% of VO_2_max) (He *et al*., 2019). Otero-Díaz et al. also proposed that bigger exercise stimuli might be needed to see an effect in humans (Otero-Díaz *et al*., 2018).

In this study, we aimed to discover possible bi-directional interactions between skeletal muscle and thermogenic adipose tissue during exercise through EVs and microRNAs in humans. Our results indicate that treatment of subcutaneous hWAs with post-exercise serum of endurance athletes after a maximum graded bicycle test to exhaustion, but not from sedentary controls, bodybuilders or sprinters, induces the expression of *UCP1* at the mRNA level. It is possible that higher oxidative metabolic adaptations in endurance athletes, as reflected by higher VO_2_max and a higher stimulus needed to achieve exhaustion compared to the other groups, contributes to a bigger response in the release of circulatory factors from the skeletal muscles or adipose tissue. This might explain why we observe changes at the *UCP1* mRNA level in the white adipocytes treated with the post-exercise serum from endurance athletes and not with the serum from the other groups.

Despite we found a clear pre-versus post-difference in the adipocytes treated with the serum from endurance athletes (using the same volume of serum per participant, corresponding to different amounts of EVs), there is not a pre-versus post-change in the cells treated with the serum-EVs (using the same amount of EVs per participant). We think that the isolation and subsequence normalization to the same amount of EVs per participant might have influenced these results, which could not reflect a more physiological effect as shown with the serum treatment. To further corroborate whether muscle EVs are directly responsible of the increased expression of *UCP1*, we treated hWAs with the same amount of muscle EVs before and after in vitro exercise-mimicking conditions using EPS. Though *UCP1* expression did not significantly increase in the adipocytes treated with muscle-EVs after EPS versus non-stimulated muscle-EVs, the increase in *UCP1* was higher in the muscle-EVs after EPS compared to the non-treated EVs control with PBS. Moreover, we show an increased in other browning markers such as *PGC1a* and *DIO2* in the hWAs treated with the muscle EVs after EPS versus non-stimulated muscle EVs. These results, including the serum treatment data, led us to the hypothesis that exercise might exert a higher response in the release of muscle-specific EVs containing microRNAs with browning potential, and hence the rationale of performing the small RNA-sequencing of serum EVs from endurance athletes before and after a bout of exercise to exhaustion.

To further investigate whether the cargo of the circulating EVs may be at least partially responsible for this effect, we screened for differentially expressed EV-associated microRNAs in endurance athletes before and after exercise that could potentially regulate *UCP1* transcription, but also BAT-to-muscle interactions, leading to three candidates: miR-196a-5p, miR-196b-5p and miR-342-3p. Mori et al. found that miR-196a-5p was able to recruit brown-like adipocytes in murine inguinal WAT through repressing its target *Hoxc8*, a WAT-specific Hox gene. Repressing *Hoxc8* led to de-repression of *C/EBPβ*, a master regulator of brown adipogenesis (Mori *et al*., 2012b), thereby increasing *Ucp1*. The expression of miR-196a-5p was increased specifically in white pre-adipocytes after cold or β-adrenergic stimulation in vivo and its expression is induced during in vitro adipogenesis. A fat-specific miR-196a-5p gain-of-function murine model, driven by the FABP4 promoter, showed higher energy expenditure and resistance to high fat diet-induced obesity (Mori *et al*., 2012b). Here, we show that miR-196a-5p is released from human skeletal myotubes EVs after EPS and after cAMP treatment. Further studies in human and murine models are needed to corroborate whether miR-196a-5p is released into the circulation from skeletal muscles acting as a browning agent of WAT. Interestingly, miR-196a-5p overexpression significantly increased *NR4A3* mRNA levels in hWAs, suggesting that miR-196a-5p may not only activate the classical *Hoxc8*/*C/EBPβ*/*Ucp1* browning axis, but also promote a broader exercise-associated metabolic transcriptional response through NR4A3. The increased trend of miR-196b-5p in muscle-derived EVs after EPS and cAMP suggests that this microRNA may be also derived from muscle cells. This is particularly interesting since miR-196b-5p share the same seed sequence as miR-196a-5p and it is also predicted to target *HOXC8* according to miR:targets prediction tools (e.g. https://www.targetscan.org). Therefore, miR-196b-5p represents a good candidate to explore in future studies taking into consideration its higher expression in serum-derived EVs from endurance athletes compared to miR-196a-5p.

Since miR-342-3p was not upregulated in muscle-EVs after in vitro stimulation, we performed EVs proteomics and integration enrichment analyses to explore protein signatures that could bring some insights of its possible cellular origin (Chen *et al*., 2022). Our data showed a strong correlation between circulating levels of EV-derived miR-342-3p and FABP4 EV protein, indicating that miR-342-3p might derive from adipocytes (Zhou *et al*., 2021). Further investigations using EV immunoprecipitation should study whether FABP4+ EVs are particularly enriched in miR-342-3p. We also found that overexpression of miR-342-3p increases the expression of *MYOG* in murine (data not shown) and hSKMs, implying a role in muscle cell differentiation. Of note, only miR-342-3p, but not miR-196a-5p nor miR-196b-5p, was enriched in white and brown adipocytes-derived EVs after cAMP treatment, indicating that adipocytes specifically release microRNA-342-3p under cAMP stimulation. Together, these results suggest that adrenergic stimulation release of EVs enriched in miR-342-3p from white and brown adipocytes may serve as a way of fat to skeletal muscle communication to promote muscle differentiation.

It is worth mentioning that the regulation of *UCP1* mRNA levels in the serum/EV-treated hWAs could be also attributed to the differential expression of other microRNAs not studied herein, but also to catecholamines, muscle-released cytokines (irisin, IL-6, meteorin-like, β-aminoisobutyric acid or BAIBA) or hepatokines (FGF-21, follistatin) that may be present in the serum and serum-derived EVs (Gonzalez-Gil & Elizondo-Montemayor, 2020). For instance, downregulation of EV-derived microRNA-27a-3p was previously proposed by Wang et al. to mediate exercise-induced browning through de-repression of PPAR-γ in obese mice (Wang *et al*., 2022). Importantly, we also detected a decreased expression of miR-27a-3p in serum-EVs from endurance athletes after exercise, and muscle-derived EVs showed a trend for decreased miR-27a expression after EPS (data not shown). To our knowledge, this is the first study in humans investigating potential muscle EV-associated microRNAs that regulate the transcriptional expression of thermogenic markers in human white adipocytes.

## Limitations

A major limitation is that adipocyte treatment readouts were transcriptional changes of *UCP1* mRNA after serum-EV and muscle-EV treatment in hWAs. This is because unlike rodent models, hWAs display very low expression of UCP1 both at the mRNA and protein levels resulting in a technical limitation in our study. This restrains the possibility of making direct comparisons between mice and humans, and the inability to determine whether increased *UCP1* transcription truly indicates protein activation and increased non-shivering thermogenesis in hWAs. In addition, due to the limiting starting volume of the human serum, this study used a very limited number of samples for the treatment of hWAs. More studies are needed to fully corroborate whether the changes at the *UCP1* mRNA levels are exclusively explained by differences in the microRNAs profile of endurance athletes or if these changes are also applicable to other training background profiles. However, we used this group to screen out for exercise-regulated microRNAs with potential browning effects.

Another limitation to acknowledge is the use of different approaches for the isolation of serum-derived EVs for small RNA sequencing (TEI) and for proteomics (ultracentrifugation). While ultracentrifugation is one of the most used procedures for EVs concentration and might constitute a purer population of exosomes, commercialized kits like polymer-based components might precipitate contaminants such a polyethylene glycol (PEG)or co-precipitate a small population of ectosomes. We particularly used TEI for small-RNA sequencing to maximize the amount of small EVs isolated from our limiting human serum sampling and to avoid substantial losses during ultracentrifugation, which usually results in lower EV yields. Similarly, we used TEI for cell treatment since high sheer stress during ultracentrifugation can negatively influence their biological functionality (Linares *et al*., 2015; Mol *et al*., 2017). Despite their convenience, polymer-based commercial kits are often not directly compatible or convenient for proteomics down-stream analyses due to the enrichment of polymers such as PEG, why we used ultracentrifugation for this approach. Yet, we implemented an optimized protocol for human serum-derived EVs proteomics using as little as 100uL of input volume. Both EVs isolated with TEI and ultracentrifugation presented EVs markers such as CD63, CD9 and CD81. Of note, we appreciated that the median size detected by NTA of the EVs isolated by TEI was slightly lower (∼100nM) than ultracentrifugation (∼150nM), which might indicate differences in EVs population. Despite these differences, the protein composition as well as the miRNA cargo of the serum-derived EVs may still retain information related to their cellular origin and hence their use for correlation analyses in this study.

Although miR-196a-5p and miR-342-3p overexpression experiments support functional potential, they do not prove that endogenous EV-mediated transfer of these microRNAs is necessary or sufficient for the effects observed after serum or EV treatment. Future studies using EV-microRNA depletion, uptake tracking, or EV subpopulation isolation is required to establish causality.

## Omics Data Repository

Raw and normalized files of the EV small RNA-sequencing can be found in the ArrayExpress collection in BioStudies Repository Database with the Accession Number: **E-MTAB-14628** (https://www.ebi.ac.uk/biostudies/studies/E-MTAB-14628).

EV mass spectrometry proteomics data have been deposited to the ProteomeXchange Consortium (http://proteomecentral.proteomexchange.org) via the PRIDE partner repository (Perez-Riverol *et al*., 2022) with the dataset identifier **PXD058619**.

## CRediT authorship contribution statement

**Dominik Tischer**: Writing – review & editing, Writing-original draft, Visualization, Methodology, Investigation and Formal analysis. **Daniela Schranner**: Resources (human exercise study). **Sebastian Kallabis**: Investigation (proteomics). **Alexander Braunsperger**: Writing – review & editing. **Martin Schönfelder**: Resources (human exercise study). **Thorsten Gnad**: Investigation (preliminary data). **Paul Jonas Jost**: Investigation (RNA-sequencing bioinformatics). **Svetozar Nesic**: Investigation (RNA-sequencing bioinformatics). **Florian Renziehausen**: Investigation (electron microscopy). **Lars Fester**: Investigation (electron microscopy). **Andreas Buness**: Supervision (RNA-sequencing bioinformatics). **Jan Hasenauer**: Supervision (RNA-sequencing bioinformatics). **Felix Meissner**: Supervision (proteomics). **Alexander Pfeifer**: Writing – review & editing, Supervision, Resources, Funding acquisition, Conceptualization. **Henning Wackerhage**: Writing – review & editing, Supervision, Resources, Funding acquisition, Conceptualization. **Ana Soriano-Arroquia**: Writing – review & editing, Writing – original draft, Visualization, Validation, Project supervision, Project administration, Methodology, Investigation, Funding acquisition, Formal analysis, Data curation, Conceptualization.

## Declaration of interests

The authors declare no competing interests.

AI was not used for the writing of this manuscript.

## Acknowledgments

This work was funded by the Deutsche Forschungsgemeinschaft (DFG, German Research Foundation) – TRR333/1 – project number 450149205 Transregio collaborative research center BATenergy to A.S.-A. and H.W. D.S was supported with a doctoral scholarship of the German National Scholarship Foundation (Studienstiftung des Deutschen Volkes). We are very grateful to Jana M. Klinke and Sophie Ameén for their excellent technical help, and Mara Neue for the culture of human brown adipocytes. We would like to thank Timo Wadenpohl and Prof. Dr. Stephanie Jung for the use of the Zetaview system. We would like to thank the Microscopy Core Facility of the Medical Faculty at the University of Bonn for providing support and instrumentation funded by the Deutsche Forschungsgemeinschaft (DFG, German Research Foundation) – project number 388171357 and the Core Unit for Bioinformatics Data Analysis of the Medical Faculty at the University of Bonn for its support in bioinformatics. We thank Unchained Labs, Royston, United Kingdom for the ExoView analyses. Graphical abstract was created using BioRender.com. Venn diagrams were generated using BioTools.fr (https://www.biotools.fr/misc/venny). Statistical analyses were performed using GraphPad Prism version 6 for Windows, GraphPad Software, Boston, Massachusetts USA, www.graphpad.com.

A pre-print version of this manuscript has been previously published in bioRxiv 2025.05.28.656129; doi: https://doi.org/10.1101/2025.05.28.656129 (Tischer *et al*., 2025).

## Supplementary information

Suppl_data_1_Database_collection

Suppl_data_2_Venn_diagram

Suppl_data_3_EV_proteomics

Suppl_data_4_up_ miRs_vs_proteom

Suppl_data_5_Figures_and_tables

