## Supplementary material for "Exercise-responsive extracellular vesicles-associated miR-196a-5p and miR-342-3p as candidate mediators of skeletal muscle-adipose tissue crosstalk": Suppl_data_5_Figures_and_tables.docx

| miR_ID | baseMean | log2FoldChange | lfcSE | padj | Levels after exercise in endurance athletes |
| --- | --- | --- | --- | --- | --- |
| hsa-let-7c-5p | 634.0771 | 0.959283 | 0.226741 | 0.000722 | Up |
| hsa-miR-140-5p | 41.29223 | 0.784401 | 0.208936 | 0.003234 | Up |
| hsa-miR-320c | 136.6493 | 0.605182 | 0.134606 | 0.000322 | Up |
| hsa-miR-155-5p | 162.6835 | 0.581794 | 0.157879 | 0.003866 | Up |
| hsa-miR-150-5p | 1382.228 | 0.568629 | 0.139828 | 0.001267 | Up |
| hsa-miR-4732-5p | 86.2688 | 0.534911 | 0.173908 | 0.021692 | Up |
| hsa-miR-181b-5p | 39.9436 | 0.529112 | 0.187672 | 0.038916 | Up |
| hsa-miR-423-5p | 3087.402 | 0.510108 | 0.082246 | 5.18E-08 | Up |
| hsa-let-7b-5p | 20139.52 | 0.501384 | 0.079007 | 4.11E-08 | Up |
| hsa-miR-183-5p | 136.8675 | 0.484952 | 0.13409 | 0.004626 | Up |
| hsa-miR-196b-5p | 64.65937 | 0.464913 | 0.159392 | 0.03289 | Up |
| hsa-miR-143-3p | 449.0068 | 0.415003 | 0.133477 | 0.021309 | Up |
| hsa-let-7a-5p | 24095.31 | 0.383317 | 0.089394 | 0.000671 | Up |
| hsa-miR-182-5p | 414.1898 | 0.344256 | 0.115334 | 0.027773 | Up |
| hsa-miR-148a-3p | 1972.494 | 0.3275 | 0.084286 | 0.00211 | Up |
| hsa-miR-342-3p | 1653.957 | 0.244928 | 0.089503 | 0.047041 | Up |
| hsa-miR-142-5p | 913.8964 | -0.24493 | 0.079057 | 0.021309 | Down |
| hsa-miR-146a-5p | 4079.762 | -0.28778 | 0.08588 | 0.010701 | Down |
| hsa-miR-93-5p | 3318.806 | -0.30255 | 0.097457 | 0.021309 | Down |
| hsa-miR-27a-3p | 193.0674 | -0.3232 | 0.119141 | 0.047739 | Down |
| hsa-miR-335-5p | 182.1853 | -0.33622 | 0.116692 | 0.035086 | Down |
| hsa-miR-148b-3p | 716.592 | -0.44554 | 0.111171 | 0.001426 | Down |
| hsa-miR-101-3p | 2079.893 | -0.50097 | 0.094889 | 8.03E-06 | Down |
| hsa-miR-199a-5p | 55.54465 | -0.58103 | 0.202948 | 0.035486 | Down |
| hsa-miR-381-3p | 35.35927 | -0.61848 | 0.226502 | 0.047041 | Down |
| hsa-miR-301a-3p | 28.84977 | -0.75924 | 0.221442 | 0.00868 | Down |

**Suppl. Table 1.** List of differentially expressed circulating exomiRs from endurance athletes after acute exercise until exhaustion.

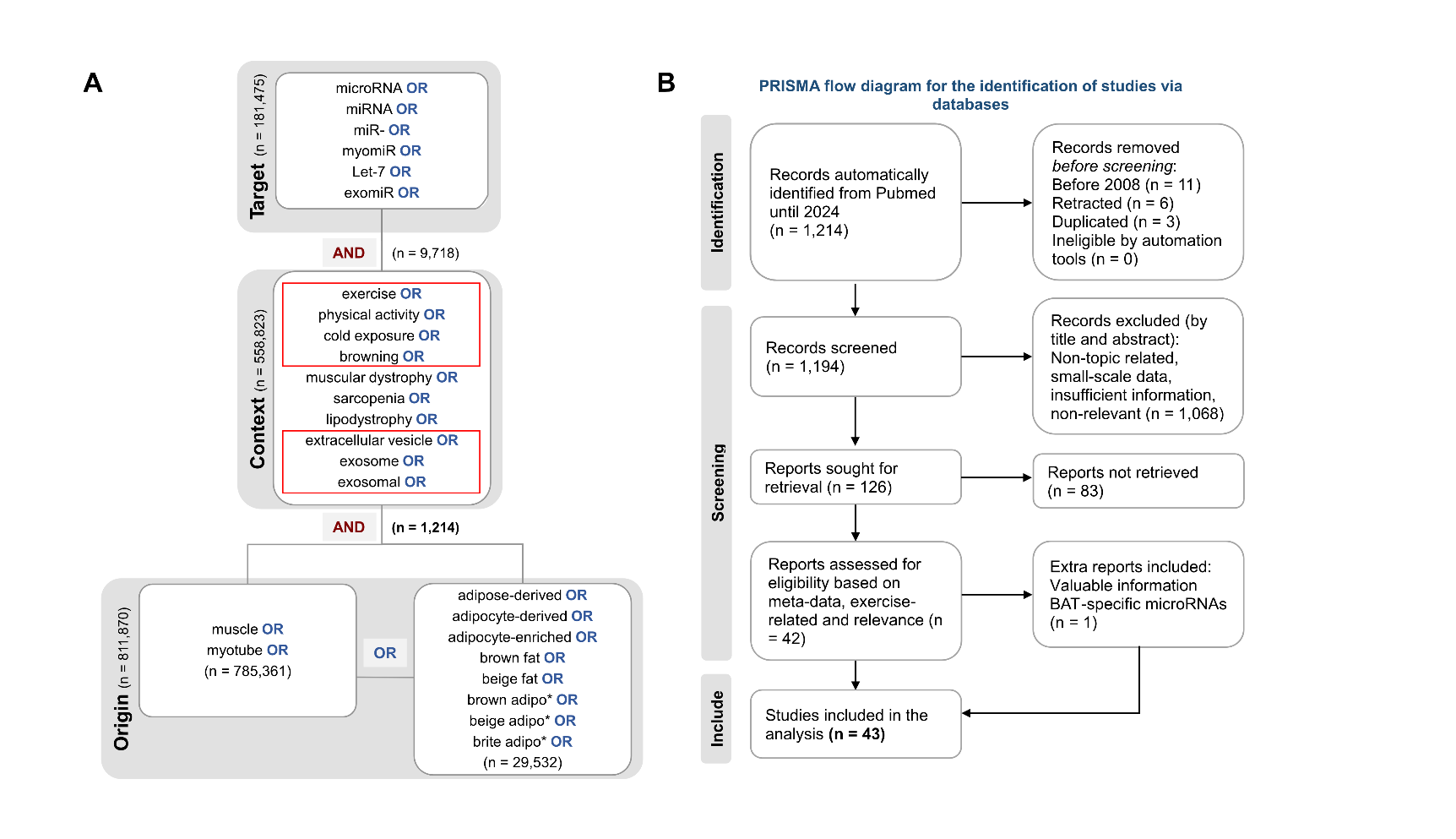

**Suppl. Fig 1.** Mining strategy used for the generation of the microRNAs and exomiRs database using NCBI published studies (A). Flow diagram used for literature screening and filtering based on PRISMA guidelines and prioritizing microarrays and small-RNA sequencing data (Page et al. 2021) (B).

| **Gene** | **Specie** | **Sequence (5’ – 3’)** | |
| --- | --- | --- | --- |
| *TBP* | Human | Forward | CACGAACCACGGCACTGATT |
|  |  | Reverse | TTTTCTTGCTGCGAGTCTGGA |
| *UCP1* | Human | Forward | AGTTCCTCACCGCAGGGAAAGA |
|  |  | Reverse | GTAGCGAGGTTTGATTCCGTGG |
| *PGC1α* | Human | Forward | CTGTGTCACCACCAAAATCCT |
|  |  | Reverse | TGTGTCGAGAAAAGGACCTTG |
| *PPARγ* | Human | Forward | GCTGTGCAGGAGATCACAGA |
|  |  | Reverse | GGGCTCCATAAAGTCACCAA |
| *DIO2* | Human | Forward | TTGAGCCGCTCCAAGTCCACTC |
|  |  | Reverse | CTGTACTGGAGACATGCACCAC |
| *CIDEA* | Human | Forward | TCAGACCTTGGGAGACAACACG |
|  |  | Reverse | CGAAGGTGACTCTCGCTATTCC |
| *NR4A3* | Human | Forward | ACTGCCCAGTAGACAAGAGACG |
|  |  | Reverse | GTTTGGAAGGCAGACGACCTCT |
| *HOXC8* | Human | Forward | AACTCGTCTCCCAGCCTCATGT |
|  |  | Reverse | TCTAGTTCCAAGGTCTGATACCG |
| *C/EBPβ* | Human | Forward | AGAAGACCGTGGACAAGCACAG |
|  |  | Reverse | CTCCAGGACCTTGTGCTGCGT |
| *MYOD1* | Human | Forward | CTCCAACTGCTCCGACGGCAT |
|  |  | Reverse | ACAGGCAGTCTAGGCTCGACAC |
| *MYOG* | Human | Forward | AGTGCCATCCAGTACATCGAGC |
|  |  | Reverse | AGGCGCTGTGAGAGCTGCATTC |

**Suppl. Table 2.** Primers sequences used for Real-Time qPCR.

| **Taqman Assay** | **Catalogue number** |
| --- | --- |
| Human *TBP* Taqman Assay | Hs00427620_m1 |
| Human *UCP1* Taqman Assay | Hs01084772_m1 |
| TaqMan Advance miRNA Assay hsa-miR-320a-3p | 478594_mir |
| TaqMan Advance miRNA Assay hsa-miR-196a-5p | 478230_mir |
| TaqMan Advance miRNA Assay hsa-miR-196b-5p | 478585_mir |
| TaqMan Advance miRNA Assay hsa-miR-342-3p | 478043_mir |
| TaqMan Advance miRNA Assay hsa-miR-342-5p | 478044_mir |

**Suppl. Table 3.** TaqMan assays used for Real-Time qPCR.

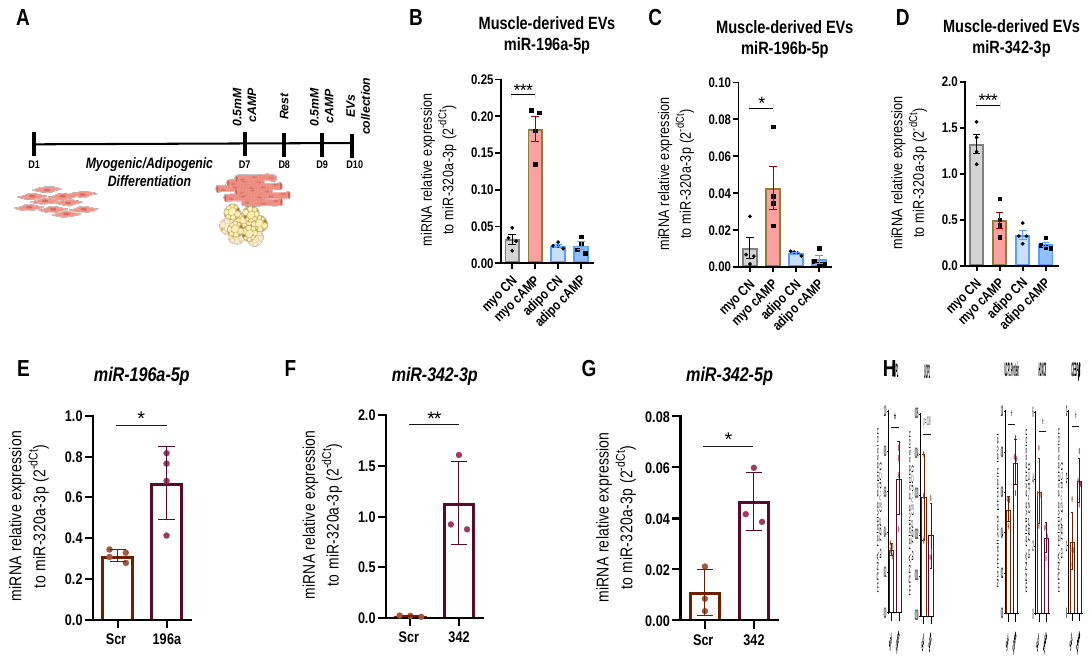

**Suppl. Fig 2.** Schematics used for cAMP treatment of murine primary muscle cells under myogenic or adipogenic differentiation (A). EVs derived from the conditioned media of myotubes but not from muscle-derived mature adipocytes are enriched in miR-196a-5p (B) and miR-196b-5p (C) after cAMP stimulation compared to the non-treated control (CN). miR-342-3p was decreased after cAMP treatment in myotubes-derived EVs (D). One-way ANOVA, Tukey’s multiple comparisons test, n = 4. Transduction validation by Real-Time qPCR: miR-196a-5p expression in miR-196 transduced hWAs (E); miR-342-3p (F) and miR-342-5p (G) expression in miR-342 transduced hWAs. *HOXC8* mRNA levels in hWAs after miR-196a overexpression (H). Two-tailed unpaired t-test compared to Scrambled control (Scr), n = 4. Error bars show SEM. * - p < 0.05; ** - p < 0.01; *** - p < 0.005.

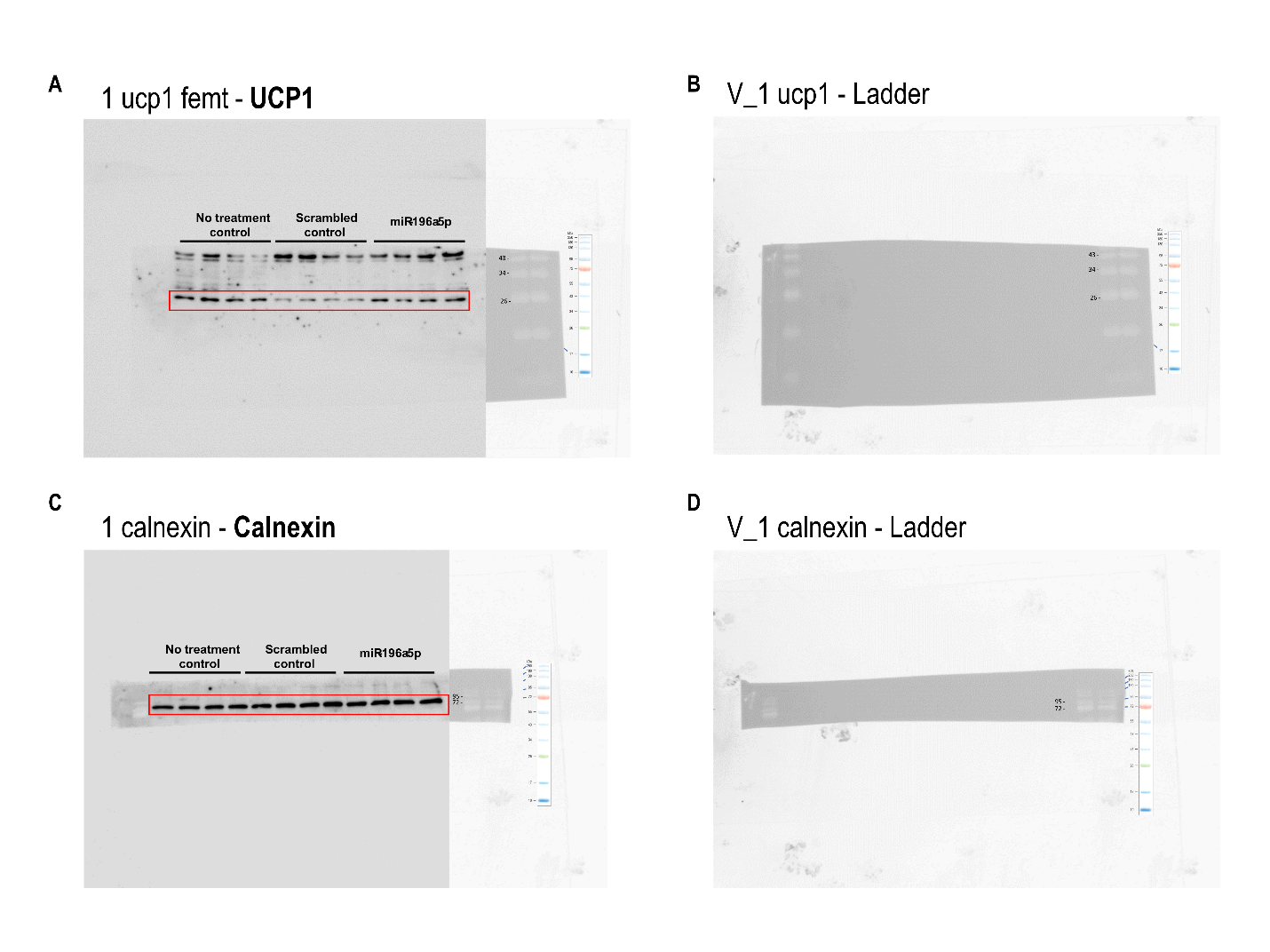

**Suppl. Fig 3.** Original western blot images for UCP1 (A, B) and calnexin (C, D) of human white adipocytes transduced with miR-196a and scrambled control.

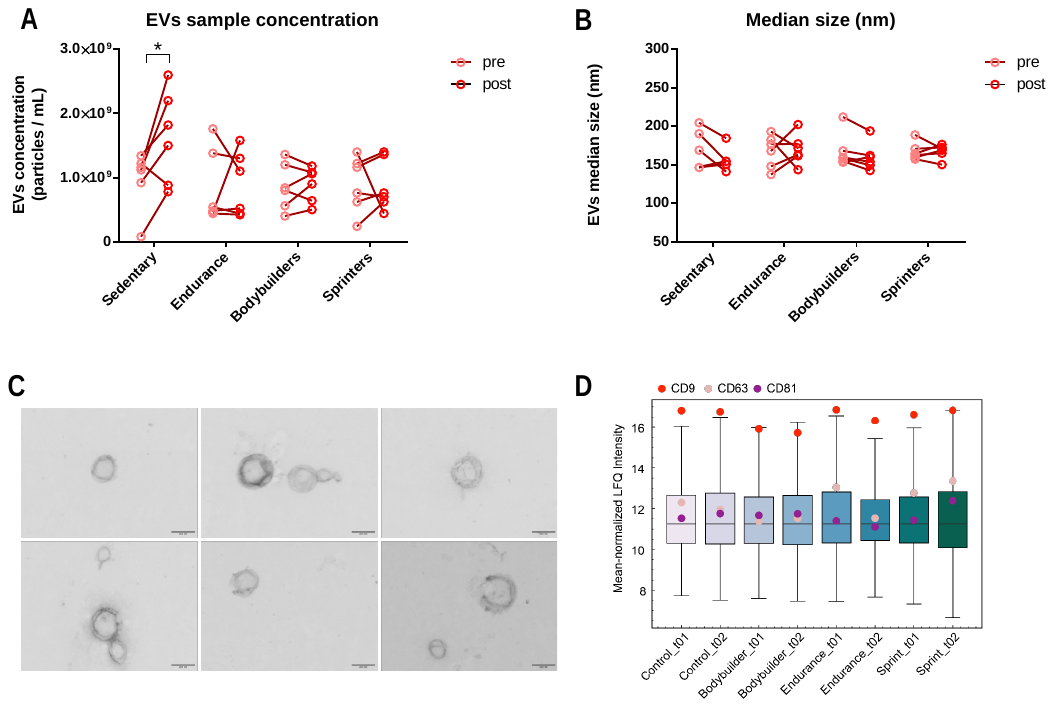

**Suppl. Fig 4.** Characterization of extracellular vesicles used for proteomics analyses. EVs sample concentration (a) and median size (b) using Nanoparticle Tracking Analysis. Two-way ANOVA, Sidak’s multiple comparisons test. n = 4-6. Error bars show SEM. * - p < 0.05; ** - p < 0.01; *** - p < 0.005. Electron microscopy images of isolated EVs (c).

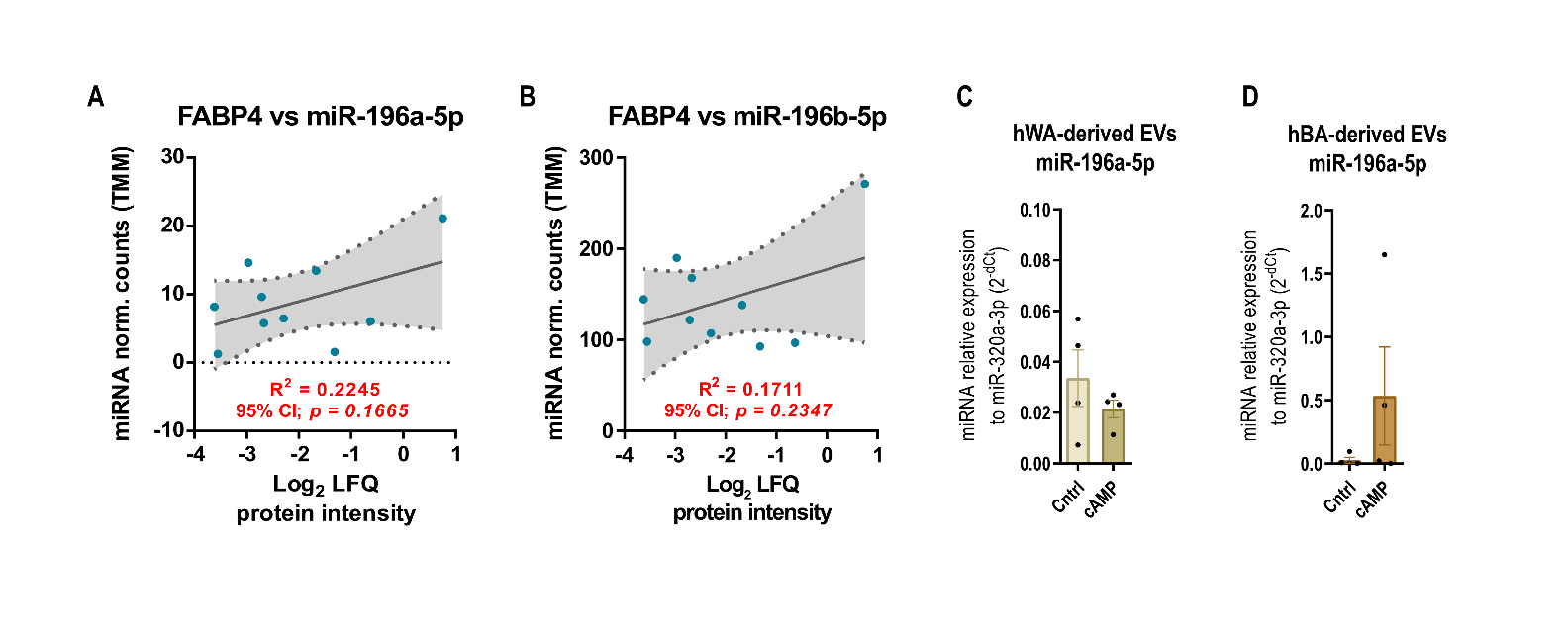

**Suppl. Fig 5.** Pearson’s correlation of miR-196a-5p (A) and miR-196b-5p (B) normalized counts versus FABP4 Log_2_ normalized protein intensity in serum-derived EVs from endurance athletes. qPCR expression of exomiRs 196a-5p in human white adipocytes (C) and human brown adipocytes (D) after cAMP treatment compared to non-treatment control. Two-tailed paired t-test. n = 4. Error bars show SEM. * - p < 0.05; ** - p < 0.01; *** - p < 0.005.
